# High-affinity P2Y_2_ and low-affinity P2X_7_ receptor interaction modulates ATP-mediated calcium signaling in murine osteoblasts

**DOI:** 10.1101/2021.03.16.435624

**Authors:** Nicholas Mikolajewicz, Delaney Smith, Svetlana V. Komarova, Anmar Khadra

## Abstract

1

P2 purinergic receptor family implicated in many physiological processes, including neurotransmission, mechanical adaptation and inflammation, consist of ATP-gated non-specific cation channels P2XRs and G-protein coupled receptors P2YRs. Different cells, including bone forming osteoblasts, express multiple P2 receptors; however, how P2X and P2Y receptors interact in generating cellular responses to various doses of [ATP] remains poorly understood. Using primary bone marrow and compact bone derived osteoblasts and BMP2-expressing C2C12 osteoblastic cells, we demonstrated conserved features in the P2-mediated Ca^2+^ responses to ATP, including a transition of Ca^2+^ response signatures from transient at low [ATP] to oscillatory at moderate [ATP], and back to transient at high [ATP], and a non-monotonic changes in the response magnitudes which exhibited two troughs at 10^−4^ and 10^−2^ M [ATP]. We identified P2Y2 and P2X7 receptors as predominantly contributing to these responses, and constructed a mathematical model of P2Y2R-induced inositol trisphosphate (IP3) mediated Ca^2+^ release coupled to a Markov model of P2X7R dynamics to study this system. Model predictions were validated using parental and CRISPR/Cas9-generated P2Y2 and P2Y7 knockouts in osteoblastic C2C12-BMP cells. Activation of P2Y2 by progressively increasing [ATP] induced a transition from transient to oscillatory to transient Ca^2+^ responses due to the biphasic nature of IP3Rs and the interaction of SERCA pumps with IP3Rs. At high [ATP], activation of P2X7R modulated the response magnitudes through an interplay between the biphasic nature of IP_3_Rs and the desensitization kinetics of P2X7Rs. Moreover, we found that P2Y2 activity may alter the kinetics of P2X7 towards favouring naïve state activation. Finally, we demonstrated the functional consequences of lacking P2Y2 or P2X7 in osteoblast mechanitransduction. This study thus provides important insights into the biophysical mechanisms underlying ATP-dependent Ca^2+^ response signatures, which are important in mediating bone mechanoadaptation.

**Author Summary:** ATP-sensitive purinergic receptors comprise a network of cell-surface receptors that activate upon ATP binding, allowing them to transmit information in a tissue- and context-dependent manner. In bone, mechanically-stimulated osteoblasts release ATP that stimulates low- and high-affinity P2 receptors in neighboring cellular populations, inducing appropriate physiological responses. P2 receptor signaling is characterized by elevations in intracellular calcium levels. When simultaneously stimulated by their common ligand, ATP, the contribution of each P2 receptor subtype gives rise to a complex calcium response, exhibiting oscillatory characteristics and biphasic dose-dependent behaviours. Here we used experimental and computational modeling approaches to determine the underlying dynamics of ATP-mediated calcium signaling in osteoblasts. The latter was done by developing a mathematical model that was comprised of a subset of low-(P2X7) and high-(P2Y2) affinity P2 receptors, reflecting the conserved P2 expression observed across different osteoblast models. We demonstrated that this model recapitulates experimental recordings of ATP-induced calcium signaling in osteoblasts and describes the dynamic interplay between P2Y2 and P2X7 receptors in the P2 receptor network.

## 3 Introduction

Extracellular ATP has long been implicated in diverse physiological functions (Verkhratsky & Burnstock, 2014), including neurotransmission (Burnstock & Ralevic, 2014), mechanical adaptation (Mikolajewicz, Mohammed, Morris, & Komarova, 2018) and the regulation of inflammation (Bours, Dagnelie, Giuliani, Wesselius, & Di Virgilio, 2011). Extracellular purines signal through 7 ionotropic receptors, i.e., the P2X ligand-gated nonspecific cation channels, and 8 metabotropic receptors, i.e., the P2Y G-protein coupled receptors (Burnstock, 2007).

ATP is the physiological agonist for all P2X receptors (P2XRs) as well as the P2Y2 and P2Y11 receptors (P2Y2R and P2Y11R, respectively) (Jacobson et al., 2006). Together they cover a range of extracellular ATP concentration ([ATP]) spanning six orders of magnitude (10^−8^ M to 10^−2^ M) (S. Xing, Grol, Grutter, Dixon, & Komarova, 2016). P2XRs are fast acting (∼10 ms activation), allowing the permeation of Na^+^, K^+^ and Ca^2+^ through the channel (Coddou, Yan, Obsil, Huidobro-Toro, & Stojilkovic, 2011) whereas P2YRs activate various types of secondary messengers, and thus act on a slower timescale than P2XRs (Erb & Weisman, 2012). Elevations in cytosolic free Ca^2+^ concentration ([Ca^2+^]_i_) is one of the hallmarks of ATP-induced signaling in many cell types, including bone-forming osteoblasts (Grol, Pereverzev, Sims, & Dixon, 2013; Mackay, Mikolajewicz, Komarova, & Khadra, 2016; Mikolajewicz, Sehayek, Wiseman, & Komarova, 2019; Mikolajewicz, Zimmermann, Willie, & Komarova, 2018; S. Xing et al., 2016). The mechanism by which P2XRs and P2YRs alter [Ca^2+^]_i_ differs: P2XR activation increases Ca^2+^ influx across the plasma membrane (North, 2002) while P2YR activation enhances Ca^2+^ release from the endoplasmic reticulum (ER) by stimulating the G_q_ protein signaling pathway, ultimately leading to the production of inositol triphosphate (IP_3_) and the activation of IP_3_ receptors (IP_3_Rs) (Burnstock, 2018). The ATP dose dependence of osteoblast responses to [ATP] was shown to be complex and does not have a clear plateau component, an outcome not explainable by the addition of individual receptor responses (S. Xing et al., 2016). While it was proposed that specific interactions between the high-affinity, mid-range and low-affinity P2Rs may explain the [ATP]-dependence, no mechanistic studies at the level of cellular signaling has yet been performed.

Markov models of P2X2/4/7R were previously developed to decipher the kinetics of ATP binding to these receptors and illustrate the interplay between receptor activation, priming, desensitization, internalization and deactivation (Khadra et al., 2013; Khadra et al., 2012; Mackay, Zemkova, Stojilkovic, Sherman, & Khadra, 2017; Yan et al., 2010; Yan, Khadra, Sherman, & Stojilkovic, 2011; Zemkova et al., 2015). Mathematical modeling has similarly been used to provide insights into the P2Y receptor signaling, particularly in the regulation of IP_3_R-mediated Ca^2+^ release (Fedorov, Rogachevskaja, & Kolesnikov, 2007; Lemon, Brockhausen, Li, Gibson, & Bennett, 2005). However, how P2X and P2Y receptors interact and what are their respective roles in generating cellular responses to various doses of [ATP] remains poorly understood.

In this study, we combined detailed experimental and computational studies of ATP-induced Ca^2+^ signals in primary mouse osteoblasts and BMP2-transfected C2C12 osteoblastic cells. We demonstrated the specific contributions of P2Y2 and P2X7 receptors to global Ca^2+^ responses using CRISPR/Cas9 -generated P2Y2 and P2Y7 knockouts in osteoblastic C2C12-BMP cell lines, and dissected the mechanisms of P2Y2 and P2X7 contributions to generating different patterns of oscillatory and sustained Ca^2+^ signals using mathematical modeling.

## 4. Results

### 4.1 ATP-mediated P2R Ca^2+^ responses in murine osteoblasts

ATP-stimulated P2R Ca^2+^ responses were assessed in three independent murine osteoblasts models: BMP2-transfected C2C12 osteoblastic cells (C2-OB), bone-marrow-derived osteoblasts (BM-OB), and compact-bone-derived osteoblasts (CB-OB). Osteoblasts were loaded with Ca^2+^-indicator dye Fura2, stimulated with varying doses of ATP, and changes in [Ca^2+^]_i_ were recorded using live cell fluorescent microscopy (**Fig. 1**). Qualitatively, the recorded Ca^2+^ response time-series signatures demonstrated a general trend of exhibiting transient single-peaked responses at low [ATP], multi-peaked oscillatory responses at mid-range [ATP], and relatively sustained single/multi-peaked response at high [ATP] (**Fig. 1A**). The Ca^2+^ responses at each [ATP] were analysed by quantifying several parameters, including overall response magnitudes and activation times, as well as oscillatory fractions, magnitudes, periods and peaks (see **Table S1** for definitions) (Mackay et al., 2016). Similar to previous study (S. Xing et al., 2016), we harmonized dose-response profiles across osteoblast models by first aligning the responses along the dose-axis to match troughs/peaks, followed by rescaling the responses to a [0,1] interval (**Fig. S1**). Such alignment allowed us to account for (*i*) inconsistencies in ATP solution preparations between experiments and (*ii*) varied dose-sensitivities across cell lines. Calcium responses induced by low [ATP] (<10^−7^ M) were consistently associated with low response magnitudes and slow activation kinetics (**Fig. 1A**, *left two columns*), with little to no oscillatory component (**Fig. 1B**). Increasing [ATP] further induced responses with faster activation kinetics and higher magnitudes (**Fig. 1A**, *middle two columns*). This also coincided with the emergence of high frequency oscillations (∼10-20 s periods; **Fig. 1B, Fig S1B, C**) which peaked at ∼10^−5^ M ATP stimulation. Notably, the oscillatory peak did not coincide with the peak magnitude. Instead, as cells were stimulated with higher [ATP], the oscillatory component began to diminish, exhibiting lower frequency oscillations and fewer oscillatory peaks, while response magnitude continued to increase, peaking at ∼2×10^−4^ M [ATP] (**Fig. 1A**, *right two columns*, **Fig. 1B**). For [ATP] >2×10^−4^ M, the response magnitude decreased with increasing [ATP] in all osteoblast lines. Thus, in all osteoblast models, the intracellular Ca^2+^ response to ATP shifts with increase in [ATP] from a transient with a single narrow peak, to oscillatory and back to transient with a pronounced wide peak.

**Figure 1.**
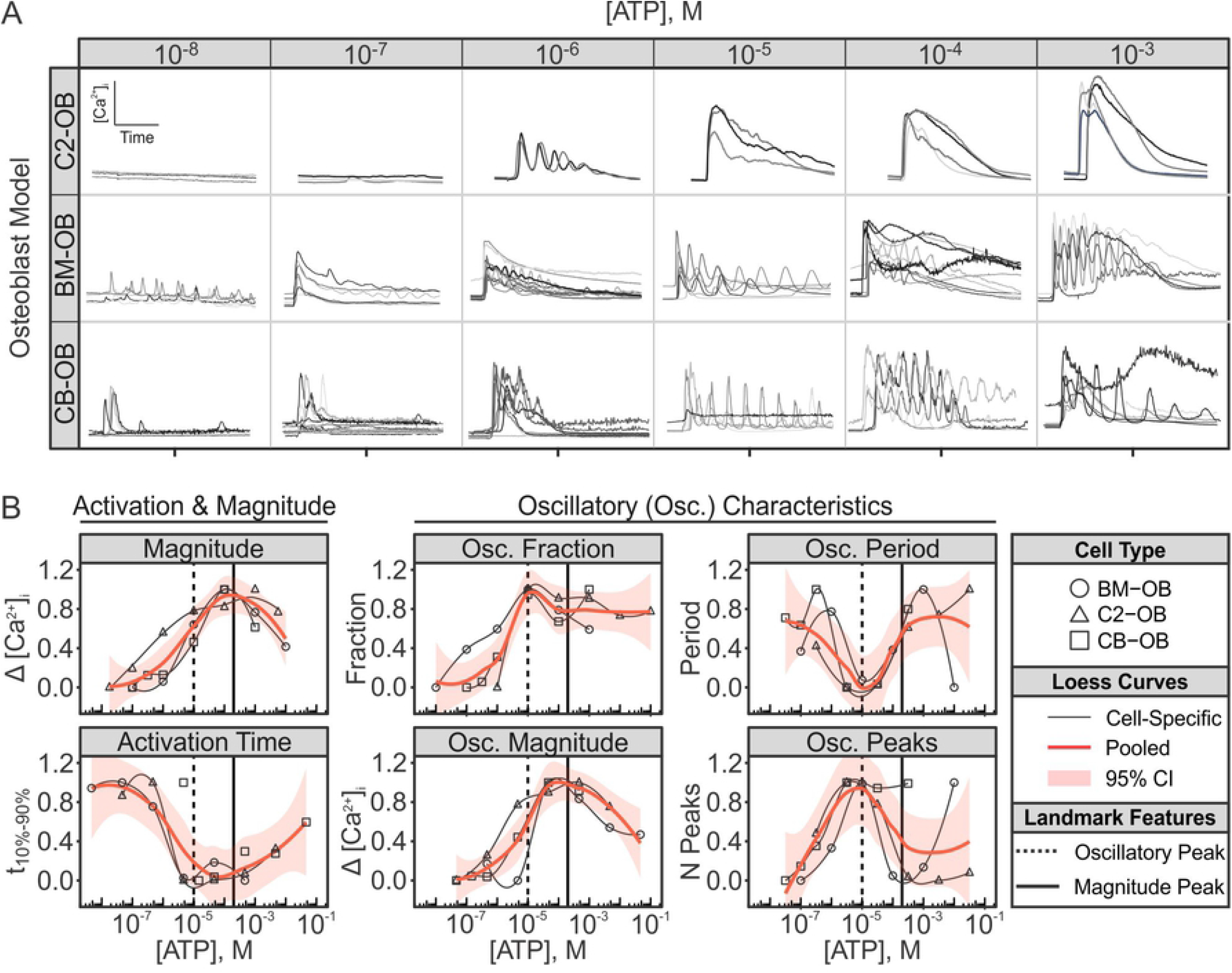
ATP-induced Ca^2+^ response characteristics are conserved across murine osteoblast lines. Fura2-loaded BMP2-transfected C2C12 osteoblastic cells (C2-OB), bone-marrow-derived osteoblasts (BM-OB), or compact-bone-derived osteoblasts (CB-OB) were stimulated with ATP (10^−8^ to 10^−3^ M), changes in [Ca^2+^]_i_ were recorded, and characteristic parameters of individual cell-level Ca^2+^ responses were quantified. (**A**) Representative ATP-induced Ca^2+^ signature responses for different osteoblastic lines. *Recording duration*: 120 s. (**B**) Activation time, magnitude and oscillatory characteristics of Ca^2+^ responses in different osteoblastic cells were aligned to obtain consensus on dose-dependency behaviours (see **Fig. S1** for intermediary alignment steps). Data are response means, normalized to peak dose-response. *Solid curves:* Loess curves fit to normalized response means. *Vertical solid lines*: peak magnitude; *Vertical dashed lines*: peak oscillatory activity. CI: confidence interval, M: molar concentration.

### 4.2 P2Y2 and P2X7 receptors orchestrate the ATP-mediated Ca^2+^ responses

To examine which P2 receptors contribute to the ATP-induced Ca^2+^ responses, we first examined their expression in osteoblastic cells of different origin. Among the P2Y family, *P2ry2, P2ry4, P2ry12*, and *P2ry14* transcripts were detected in all osteoblastic cells by RT-qPCR (**Fig. 2A** *top*). Among the P2X family, *P2rx4* and *P2rx7* transcripts were the most abundantly expressed in all osteoblastic cells (**Fig. 2A** *bottom*). These expression profiles suggest that P2Y2, 4, 12, 14 and P2X4, 7 are the predominant P2 receptor subtypes expressed in osteoblastic cells, among which P2Y2 and P2X7 were the most abundant transcripts. To confirm that P2Y2 and P2X7 receptors are functional, we stimulated Fura2-loaded C2-OB cells with ATP and receptor-specific agonists: the P2Y2-agonist UTP and P2X7-agonist bzATP (**Fig. 2B**). Consistent with previously characterized P2 receptor sensitivities (S. Xing et al., 2016), we found that the estimated EC50s were 1.0 µM for ATP, 2.8 µM for UTP and 26.4 µM for bzATP in C2-OB cells. Importantly, the oscillatory Ca^2+^ responses evoked by 10^−6^ M ATP were recapitulated following stimulation with 10^−6^ M UTP, and similarly, the sustained responses evoked by 1 mM ATP were observed following 1 mM bzATP stimulation, suggesting that P2Y2 and P2X7 receptors dominate the responses to lower and higher [ATP], respectively (**Fig. 2C**). Using CRISPR-Cas9 double-nickase constructs, we generated clonal C2-OB cells harboring mutations in *P2ry2* (*P2ry2*Δ) or *P2rx7 (P2rx7*Δ) (**Fig. 2D**) to further investigate the independent contribution of P2Y2 and P2X7 to the P2-mediated Ca^2+^ responses (presented in subsequent sections).

**Figure 2.**
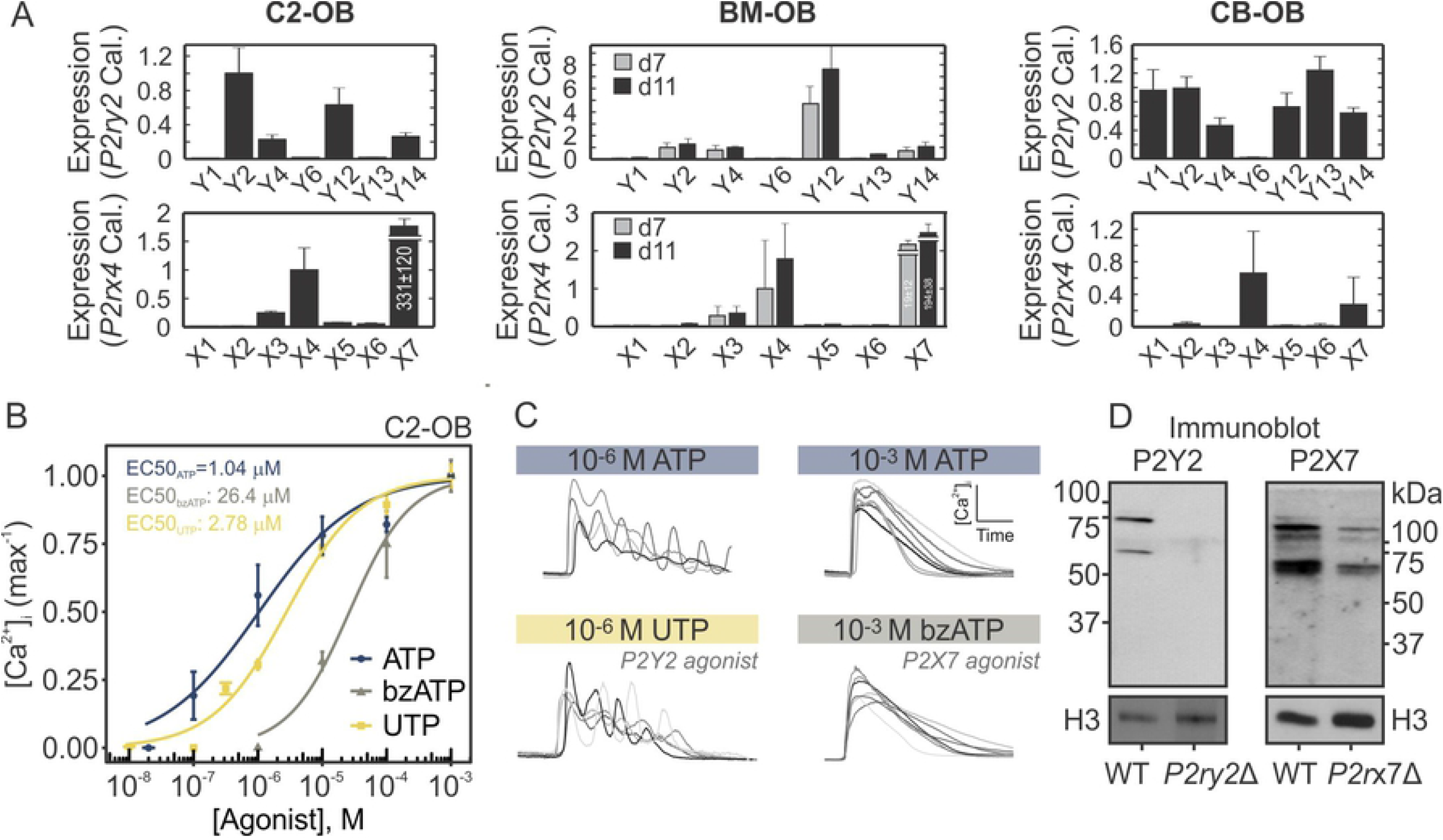
Functional P2Y2 and P2X7 are expressed in osteoblastic cells. (**A**) P2 expression determined by RT-qPCR in C2-OB, BM-OB, and CB-OB. Relative transcript expression was calculated by ΔΔCT method, and *P2ry2* and *P2rx4* were used as calibrators for P2Y and P2X receptors, respectively. Data are means ± SEM, n = 3 independent cultures per cell line. **(B)** Fura2-loaded C2-OB cells were stimulated by ATP, UTP, or BzATP and [Ca^2+^]_i_ response magnitudes were measured. Data are normalized means ± SEM (*markers*) fitted with hill functions (*curves*) for their dose-response curves. **(C)** Representative Ca^2+^ responses observed in C2-OB stimulated by 10^−6^ M ATP or UTP, and 10^−3^ M ATP or BzATP. (**D**) P2Y2 and P2X7 protein expression assessed by immunoblot in WT, *P2ry2*Δ and *P2rx7*Δ C2-OB whole cell lysates. Histone H3 was used as a loading control.

### 4.3 A flux-balance-based model of P2Y2R and P2X7R driven Ca^2+^ responses

To decipher the underlying biophysical mechanisms governing the ATP-stimulated Ca^2+^ responses in osteoblasts, we developed a mathematical model that integrates P2Y2R-mediated Ca^2+^ release, as described by the Li-Rinzel model of the IP_3_R (Y. Li & Rinzel, 1994) with a Markov model of P2X7R kinetics adapted from (Khadra et al., 2013) (**Fig. 3**). The cell was divided into two compartments (**Fig. 3**A), the endoplasmic reticulum (ER) and the cytosol, with Ca^2+^ concentrations in each compartment denoted by [Ca^2+^]_ER_ and [Ca^2+^]_i_, respectively. The detailed description of the model is given in *Methods*. Briefly, the model describes (*i*) Ca^2+^ mobilization across cell membrane, including Ca^2+^ influx through P2X7R receptor channels (*J*_*P2X7*_) and the constant inward leak (*J*_*INleak*_), as well as Ca^2+^ efflux through the plasma-membrane-Ca^2+^-ATPase (PMCA) pumps (*J*_*PMCA*_); and (*ii*) Ca^2+^ mobilization across ER membrane, including P2Y2R-mediated Ca^2+^-induced Ca^2+^-release (CICR) through IP_3_R (*J*_*IP3R*_), Ca^2+^ leak across the ER membrane (*J*_*ERleak*_), and Ca^2+^ uptake through the sarco/endoplasmic Ca^2+^ ATPase (SERCA) pumps (*J*_*SERCA*_). The reduced two-dimensional Li-Rinzel model for IP_3_R-mediated CICR was chosen for its simplicity and ability to produce transitions between the desired modes of activity; it follows the Hodgkin-Huxley gating formalism (see *Methods*), with two fast activation variables and one slow inactivation variable that depend on [IP_3_] and [Ca^2+^]_i_, producing an open probability profile for CICR that is biphasic with respect to [Ca^2+^]_i_. Given that P2Ys modulate intracellular Ca^2+^ responses indirectly by stimulating IP_3_ production, an equation describing [ATP]-dependent IP_3_ production was added to the P2Y2R submodel.

**Figure 3.**
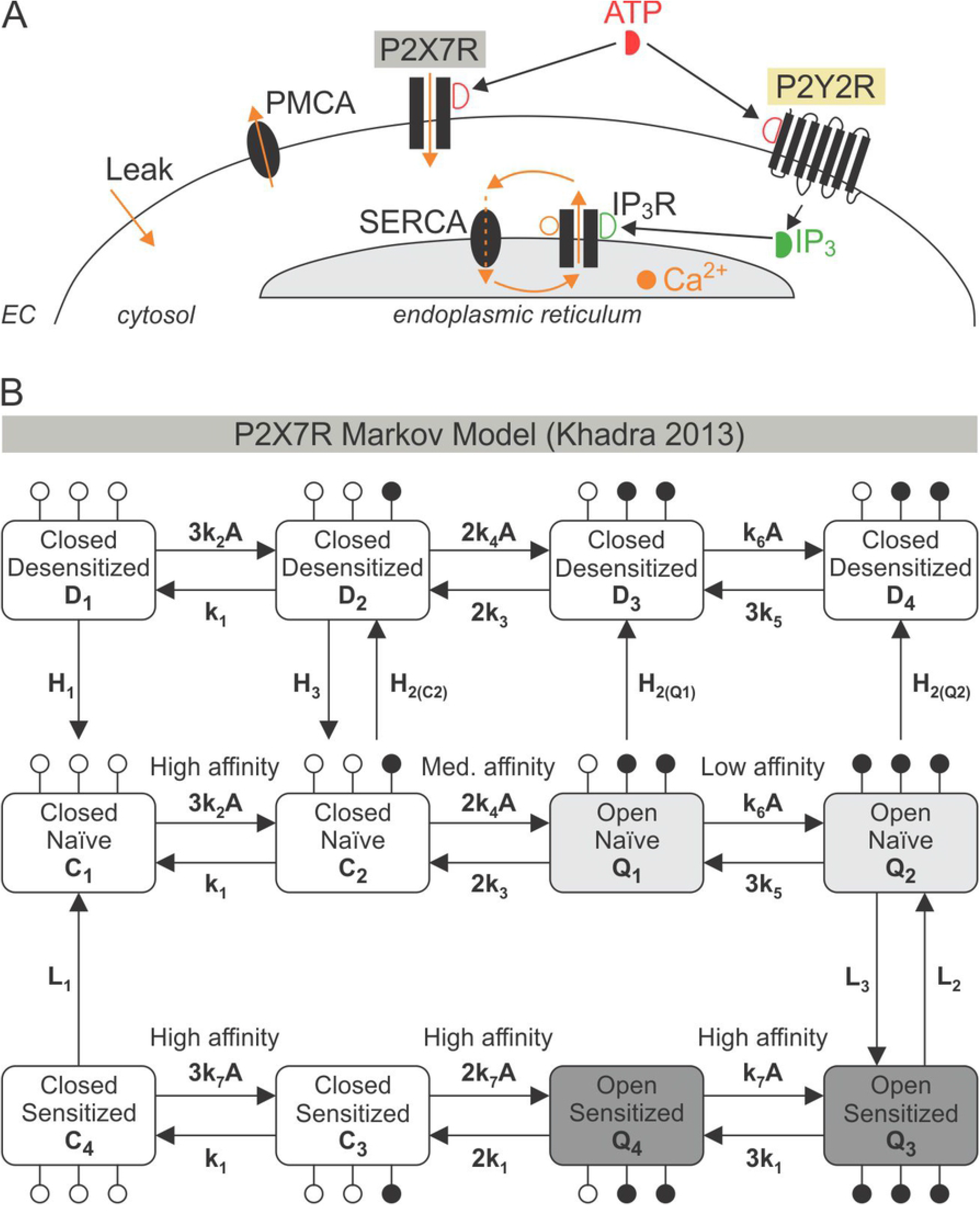
Schematic of the mathematical model describing P2 receptor-mediated Ca^2+^ responses. **(A)** ATP activates P2X7 and P2Y2 receptors on the plasma membrane, stimulating Ca^2+^ entry and IP_3_ production, respectively. IP_3_ production leads to Ca^2+^ release from the endoplasmic reticulum (ER) through IP_3_Rs. Sarco/endoplasmic Ca^2+^ ATPase (SERCA) activity replenishes the ER and allows for Ca^2+^ oscillations when combined with the biphasic response of IP_3_Rs to Ca^2+^ due to Ca^2+^-induced Ca^2+^-release (CICR). Ca^2+^ is removed from the cell by plasma membrane Ca^2+^ ATPases (PMCA). A constant inward Ca^2+^ leak ensures positive [Ca^2+^]_i_ in the absence of ATP. **(B)** Schematic of the P2X7R Markov Model. *Middle, lower and upper rows:* Fraction of P2X7Rs in naïve, sensitized and desensitized states, respectively. *Open and solid circles*: Sites unoccupied and occupied by ATP, respectively. Receptors in the closed (*C*_*i*_) and desensitized (*D*_*i*_) states have closed channel pores, whereas receptors in the open (*Q*_*i*_) states, have open channel pores with identical conductance, *i* = 1 ― 4. Model parameter values are listed in **Table 1**.

Ca^2+^ flux through P2X7R, *J*_*P2X7*_, on the other hand, was determined by the Ca^2+^ current (*I*_*P2X7*_) through the receptor channels generated by a 12-state Markov P2X7R sub-model (**Fig. 3B**) (Khadra et al., 2013). The P2X7R submodel assumes that each receptor has three ATP binding sites, two of which must be occupied for the receptor to be open, and that each state represents the fraction of receptors with a given number of occupied ATP-binding sites (**Fig. 3B**, *solid circles*). The closed, *C*_*i*_, and desensitized, *D*_*i*_, states are non-conducting, whereas the open states *Q*_*i*_ (*i* = 1 ― 4) possess the same conductance *g*_*X7*_. The states were divided into three rows corresponding to desensitized (**Fig. 3B**, *top row*), naïve (**Fig. 3B**, *middle row*) and sensitized or primed (**Fig. 3B**, *bottom row*) states, respectively. The naïve row is comprised of the states *C*_1_, *C*_2_, *Q*_1_, *Q*_2_ that have not been exposed to ATP for a prolonged period of time, whereas the sensitized and desensitized rows are comprised of the states *C*_3_, *C*_4_, *Q*_3_, *Q*_4_ (*D*_1_, *D*_2_, *D*_3_, *D*_4_) that have been previously exposed to ATP. The forward and backward transitions along each row describes ATP binding and unbinding, respectively, whereas downward and upward transitions between the rows represent receptor sensitization (middle to bottom row), desensitization (middle to top row) or recovery (bottom/top to middle row). The rate of desensitization increases as more ATP molecules bind to P2X7R and the open probability along the sensitized row is larger than that for the naïve row.

**Table 1.**
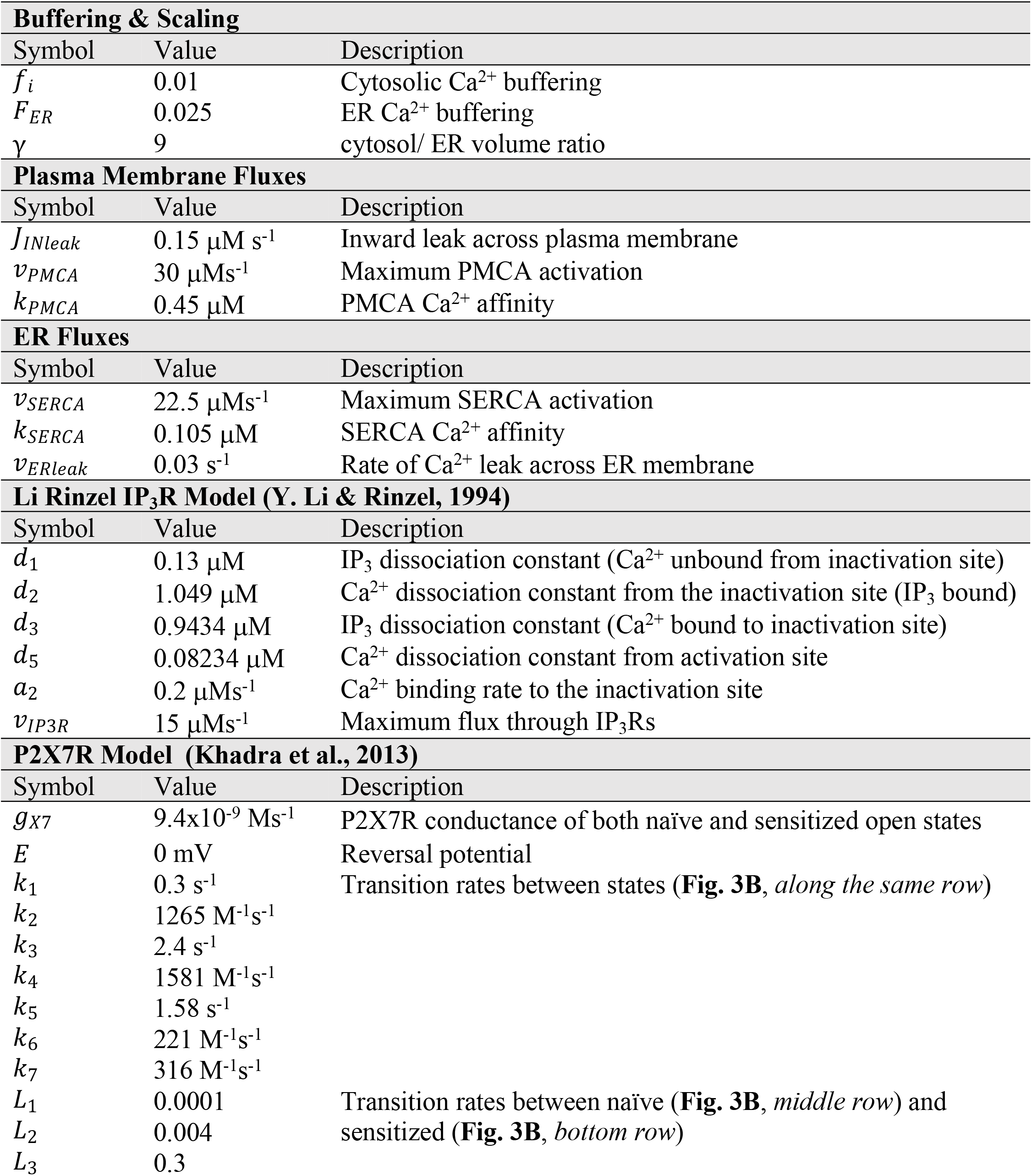
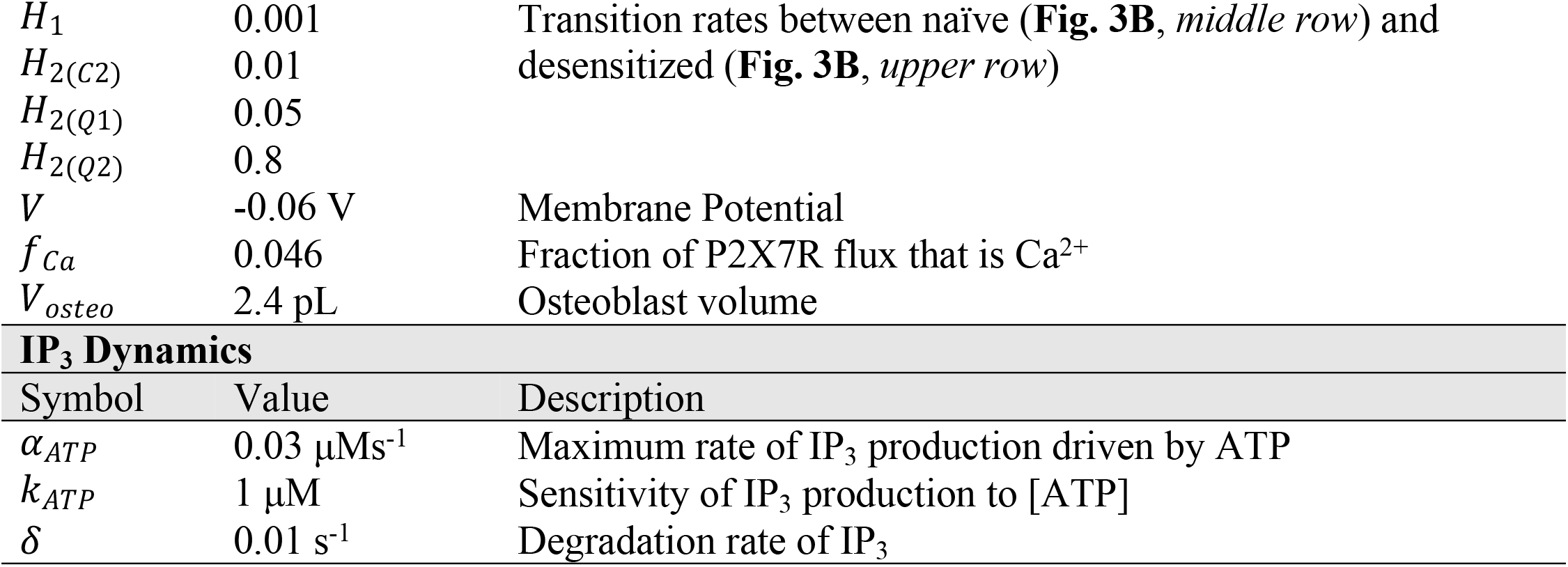
Mathematical Model Parameters

| **Table 1. Mathematical Model Parameters** |  |  |
| --- | --- | --- |
| **Buffering & Scaling** |  |  |
| Symbol | Value | Description |
| $f_i$ | 0.01 | Cytosolic $\text{Ca}^{2+}$ buffering |
| $F_{ER}$ | 0.025 | ER $\text{Ca}^{2+}$ buffering |
| $\gamma$ | 9 | cytosol/ ER volume ratio |
| **Plasma Membrane Fluxes** |  |  |
| Symbol | Value | Description |
| $J_{INleak}$ | $0.15 \mu\text{M s}^{-1}$ | Inward leak across plasma membrane |
| $v_{PMCA}$ | $30 \mu\text{Ms}^{-1}$ | Maximum PMCA activation |
| $k_{PMCA}$ | $0.45 \mu\text{M}$ | PMCA $\text{Ca}^{2+}$ affinity |
| **ER Fluxes** |  |  |
| Symbol | Value | Description |
| $v_{SERCA}$ | $22.5 \mu\text{Ms}^{-1}$ | Maximum SERCA activation |
| $k_{SERCA}$ | $0.105 \mu\text{M}$ | SERCA $\text{Ca}^{2+}$ affinity |
| $v_{ERleak}$ | $0.03 \text{ s}^{-1}$ | Rate of $\text{Ca}^{2+}$ leak across ER membrane |
| **Li Rinzel $\text{IP}_3\text{R}$ Model (Y. Li & Rinzel, 1994)** |  |  |
| Symbol | Value | Description |
| $d_1$ | $0.13 \mu\text{M}$ | $\text{IP}_3$ dissociation constant ($\text{Ca}^{2+}$ unbound from inactivation site) |
| $d_2$ | $1.049 \mu\text{M}$ | $\text{Ca}^{2+}$ dissociation constant from the inactivation site ($\text{IP}_3$ bound) |
| $d_3$ | $0.9434 \mu\text{M}$ | $\text{IP}_3$ dissociation constant ($\text{Ca}^{2+}$ bound to inactivation site) |
| $d_5$ | $0.08234 \mu\text{M}$ | $\text{Ca}^{2+}$ dissociation constant from activation site |
| $a_2$ | $0.2 \mu\text{Ms}^{-1}$ | $\text{Ca}^{2+}$ binding rate to the inactivation site |
| $v_{\text{IP}_3\text{R}}$ | $15 \mu\text{Ms}^{-1}$ | Maximum flux through $\text{IP}_3\text{Rs}$ |
| **P2X7R Model (Khadra et al., 2013)** |  |  |
| Symbol | Value | Description |
| $g_{X7}$ | $9.4 \times 10^{-9} \text{ Ms}^{-1}$ | P2X7R conductance of both naïve and sensitized open states |
| $E$ | 0 mV | Reversal potential |
| $k_1$ | $0.3 \text{ s}^{-1}$ | Transition rates between states (**Fig. 3B**, *along the same row*) |
| $k_2$ | $1265 \text{ M}^{-1}\text{s}^{-1}$ | |
| $k_3$ | $2.4 \text{ s}^{-1}$ | |
| $k_4$ | $1581 \text{ M}^{-1}\text{s}^{-1}$ | |
| $k_5$ | $1.58 \text{ s}^{-1}$ | |
| $k_6$ | $221 \text{ M}^{-1}\text{s}^{-1}$ | |
| $k_7$ | $316 \text{ M}^{-1}\text{s}^{-1}$ | |
| $L_1$ | 0.0001 | Transition rates between naïve (**Fig. 3B**, *middle row*) and sensitized (**Fig. 3B**, *bottom row*) |
| $L_2$ | 0.004 | |
| $L_3$ | 0.3 | |

| $H_1$ | 0.001 | Transition rates between naïve (**Fig. 3B**, *middle row*) and desensitized (**Fig. 3B**, *upper row*) |
| --- | --- | --- |
| $H_{2(C2)}$ | 0.01 | |
| $H_{2(Q1)}$ | 0.05 | |
| $H_{2(Q2)}$ | 0.8 | |
| $V$ | -0.06 V | Membrane Potential |
| $f_{Ca}$ | 0.046 | Fraction of P2X7R flux that is $Ca^{2+}$ |
| $V_{osteo}$ | 2.4 pL | Osteoblast volume |
| **IP<sub>3</sub> Dynamics** |  |  |
| Symbol | Value | Description |
| $\alpha_{ATP}$ | 0.03 $\mu\text{Ms}^{-1}$ | Maximum rate of IP<sub>3</sub> production driven by ATP |
| $k_{ATP}$ | 1 $\mu\text{M}$ | Sensitivity of IP<sub>3</sub> production to [ATP] |
| $\delta$ | 0.01 $\text{s}^{-1}$ | Degradation rate of IP<sub>3</sub> |

Combining the two submodels together produced the following model:

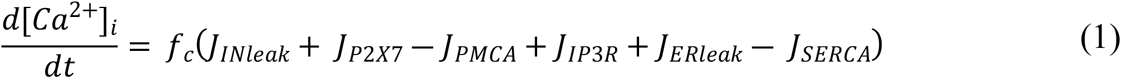

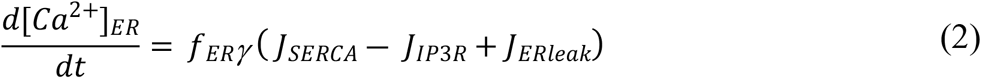

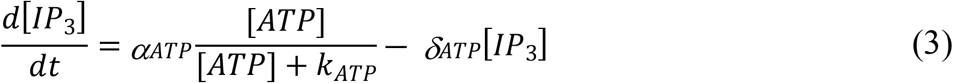

where *f*_*c*_ and *f*_*ER*_ represent the fraction of free Ca^2+^ in the cytosol and ER, respectively, as a result of buffering with 0 < *f*_*c*_ < *f*_*ER*_ << 1, *γ* is the ratio between cytosolic and ER volume, *α*_*ATP*_ is the maximum rate of IP_3_ production by P2Y2 in response to ATP, *K*_*ATP*_ is the half-maximum production of IP_3_ through P2Y2R and *δ*_*ATP*_ is the rate of IP_3_ degradation.

Using the model parameters provided in **Table 1**, we simulated Ca^2+^ responses to different [ATP] in three specific cases: (*i*) in naïve cells expressing both P2Y2R and P2X7R using the full model (**Fig. 4A**, *blue curves*), (*ii*) in cells that do not express P2Y2R, using the submodel for P2X7 component only (**Fig. 4A**, *grey curves*), and (*iii*) in cells that do not express P2X7R using the submodel for P2Y2 component only (**Fig. 4A**, *yellow curves*). The simulated Ca^2+^ responses were compared with those obtained experimentally in WT (**Fig. 4B**, *blue curves*), *P2ry2*Δ (**Fig. 4B**, *grey curves*) and *P2rx7*Δ (**Fig. 4B**, *yellow curves*) C2-OB cells. As shown, the responses to low [ATP] were predominantly P2Y2-mediated, while the response to high [ATP] were jointly mediated by P2Y2 and P2X7. Notably, the characteristic two-peaked response to 10^−3^ M observed in experimental recordings (**Fig. 1A, Fig. 4B**) was predicted by the full model (**Fig 4A**) of WT cells, but was abolished in *P2ry2*Δ and *P2rx7*Δ recordings and in simulations of P2X7 and P2Y2 submodels. These data strongly support the interaction between P2Y2 and P2X7 receptors in generating this unique signaling feature.

**Figure 4.**
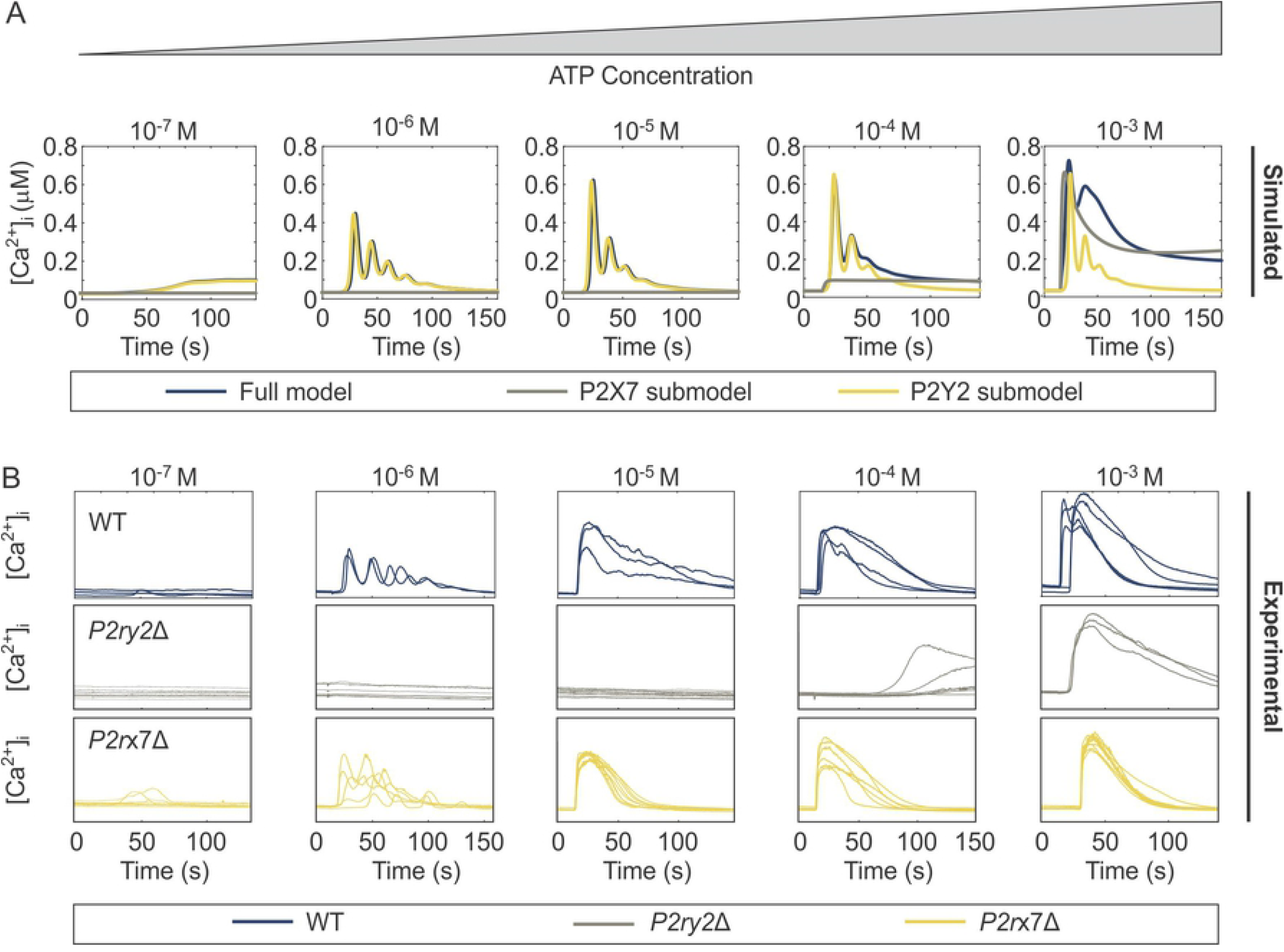
Comparison of simulated and experimental dynamics of ATP-induced [Ca^2+^]_i_ responses. **(A)** Time series simulations of [Ca^2+^]_i_ responses generated by the complete model of P2Y2 and P2X7 (full model, *blue*), P2X7 submodel (*grey*) and P2Y2 submodel (*yellow*). Parameter values are provided in **Table 1. (B)** Experimental recordings of [Ca^2+^]_i_ responses in WT (*blue*), *P2ry2*Δ (*grey*) and *P2rx7*Δ (*yellow*) C2-OB cells in response to varying [ATP].

One interesting aspect of the recordings and simulations displayed in **Fig. 4** was the observation of oscillatory Ca^2+^ responses at intermediate [ATP], with transient responses at low and high [ATP], indicating that the mathematical model developed in this study recapitulates the characteristic Ca^2+^ signatures observed in C2-OB cells over a physiological range of [ATP].

### 4.4 P2Y2 drives the transition from transient to oscillatory Ca^2+^ responses

Since the oscillatory component required P2Y2 activity in both experimental and simulated responses to ATP, we next investigated how the transition between transient and oscillatory responses is achieved by the P2Y2 receptor. Given that [IP_3_] and [Ca^2+^]_ER_ are slow variables in the model defined by **Eqs. (1)-(3)**, we set *J*_*P2X7R*_ = 0 and applied slow-fast analysis on the resulting P2Y2 receptor model by assuming that these two variables change slowly relative to other “fast” variables in the model. We set the two variables ([IP_3_] and [Ca^2+^]_ER_) as independent adjustable parameters in the P2Y2 model, and investigated how the steady state dynamics of fast variables change when [IP_3_] and [Ca^2+^]_ER_ are altered. The two-parameter bifurcation diagram (**Fig 5**) exhibited an oscillatory region in the “parameter” space formed by [IP_3_] and [Ca^2+^]_ER_ within which [Ca^2+^]_i_ is periodic (**Fig. 5**, *grey region*). Outside this region, [Ca^2+^]_i_ attained steady state values that are [IP_3_] and [Ca^2+^]_ER_-dependent. The boundary of this oscillatory region (generated by two supercritical Hopf bifurcations) defined the threshold for [Ca^2+^]_i_ to transition between these two main patterns of activity: quiescence and oscillatory. Thus the application of increasing [ATP] in this model would induce an increase in IP_3_ (since [*IP*_3_] ∝ [*ATP*]) and a decrease in [Ca^2+^]_ER_, generating three possible scenarios for the time courses of [Ca^2+^]_i_. When low [ATP] is applied the trajectory stays to the left of the oscillatory region (**Fig. 5A**, *red arrow*) because the ATP-induced IP3 increase is low (**Fig. 5B**, *red curve*) and [Ca^2+^]_ER_ remains high (**Fig. 5C**, *red curve*), resulting in a low magnitude persistent transient response **(Fig. 5D**, *red curve*). Intermediate [ATP] results in a hybrid response that becomes periodic when the trajectory crosses the left boundary of the oscillatory regime (**Fig. 5A**, *green curve*), due to a higher increase in IP3 and a faster decrease in [Ca^2+^]_ER_ (**Fig. 5 B**,**C**, *green curves*), leading to a response characterized by damped oscillations (**Fig. 5D**, *green curve*). When high [ATP] is applied, the trajectory crosses the oscillatory region very briefly (**Fig. 5A**, *blue curve*) because even though the ATP-induced IP3 increase is higher (**Fig. 5B**, *blue curve*), the [Ca^2+^]_ER_ does not decrease as fast (**Fig. 5 C**, *blue curve*) due to the biphasic nature of IP3Rs incorporated in the P2Y2 sub-model and the interaction of SERCA pumps with IP3Rs (Y. Li & Rinzel, 1994); this results in a high magnitude semi-persistent response (**Fig. 5D**, *blue curve*). The aforementioned mechanism suggests that the heterogeneity in Ca^2+^ response profiles at a given [ATP] observed experimentally (**Fig. 1A**) may be due to variations in the initial conditions, such as the expression levels of PMCA/SERCA pumps and IP_3_R, potentially leading to differences in the initial [IP_3_] and [Ca^2+^]_ER_. These simulations demonstrate that the oscillatory response strongly depends on the initial [IP_3_] and [Ca^2+^]_ER_, and that when increasing doses of [ATP] are applied, P2Y2-mediated changes in [IP_3_] and [Ca^2+^]_ER_ result in different patterns of Ca^2+^ responses.

**Figure 5.**
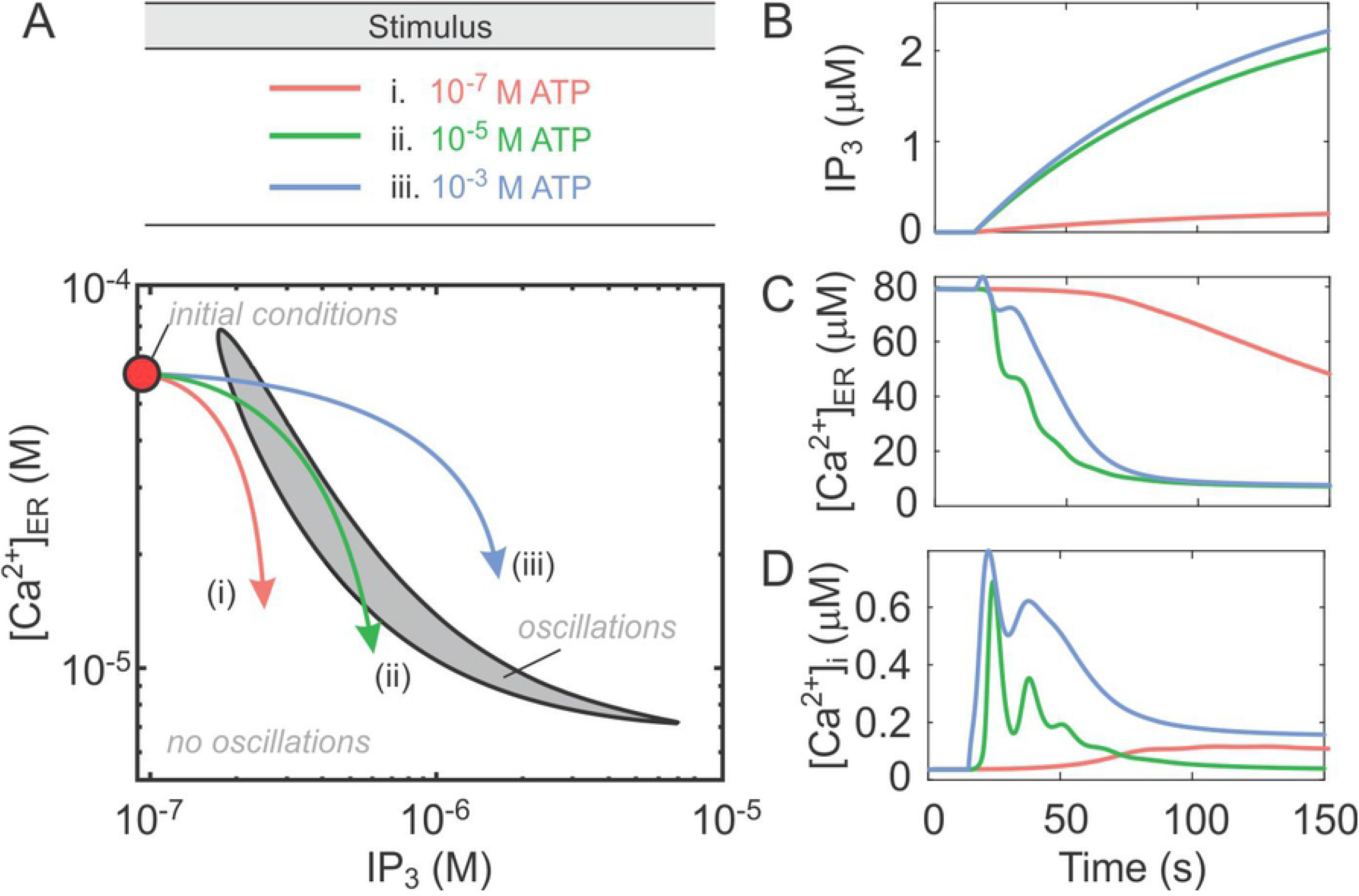
Ca^2+^ oscillatory response dynamics defined by the P2Y2-induced changes in [Ca^2+^]_ER_ and [IP_3_]. The model defined by **Eqs. (1)-(3)**, was examined with *J*_*P2X7R*_ = 0. **(A)** The slow variables representing [IP3] and [Ca^2+^]_ER_ were set to be independent parameters and the continuation method in XPPATU was applied to track two supercritical Hopf bifurcation points that enclose the oscillatory region (*grey region*). *Arrows* indicate the three possible scenarios that describe the changes in [IP_3_] and [Ca^2+^]_ER_ during Ca^2+^ responses: 10^−7^ M ATP response trajectory remains outside the oscillatory region (*red*), 10^−5^ M ATP response trajectory spends an extended period of time inside the oscillatory region (*green*) and 10^−3^ M ATP response trajectory briefly crosses through the oscillatory region (*blue*). **(B-D)** Simulated changes in IP3 **(B)**, [Ca^2+^]_ER_ **(C)** and [Ca^2+^]_i_ **(D)** following the application of 10^−7^ M ATP (*red*) 10^−5^ M ATP (*green*) or 10^−3^ M ATP (*blue*).

### 4.5 P2X7 modulates the magnitude of Ca^2+^ response to ATP

We next investigated the non-monotonic [ATP]-dependent dose-response profile for the magnitude of Ca^2+^ response observed across all osteoblastic lines (**Fig. 1B**). Using the model defined by **Eqs. (1)-(3)**, we generated Ca^2+^ responses to [ATP] ranging between 10^−8^ to 10^−2^ M ATP (with 10^−8^ M [ATP] increments), and computed the maximum [Ca^2+^]_i_ reached within 120 s (**Fig. 6A**). The full model (**Fig. 6A**, *blue curve*) recapitulated the experimental magnitude dose-response profile (**Fig. 6A** *blue triangles*), including the troughs at moderate (**Fig. 6A**, *light grey region*) and elevated (**Fig. 6A**, *dark grey region*) [ATP] within the plateau component of the response. This dose-dependency became monotonic in experimental recordings of Fura2-loaded *P2rx7*Δ C2-OB cells stimulated with ATP (**Fig. 6A**, *red circles*) and in the model lacking P2X7Rs (**Fig. 6A**, *red curve*), strongly implicating P2X7 in this phenomenon. Therefore, we next plotted the total Ca^2+^ flux through P2X7Rs, estimated as the area under the P2X7R flux curve, as well as the maximal Ca^2+^ fluxes through P2X7Rs and IP_3_Rs predicted by the model (**Fig. 6B-E**). When P2X7R-mediated Ca^2+^ entry became evident at 10^−5^ M ATP (**Fig. 6B**, *light grey region*), the maximum flux through IP_3_Rs in the full model (**Fig. 6C**, *black curve*) dropped below that of the P2X7R-lacking submodel (**Fig. 6C**, *red curve*) due to the biphasic dependence of IP_3_Rs on [Ca^2+^]_i_, resulting in the first trough in the dose-response (**Fig. 6A**, *light grey region*). At high 10^−2^ M [ATP], on the other hand, the time required for the P2X7R flux to decay to half of its maximum (t_1/2_) decreased (**Fig. 6D**). As a result, despite the maximum P2X7R flux continually increasing (**Fig. 6E**), the Ca^2+^ entering through P2X7R began to decrease at elevated [ATP] (**Fig. 6B**, *dark gre*y), resulting in the second trough in the dose-response (**Fig. 6A**, *dark grey*). Taken together, these simulations indicate that the non-monotonic Ca^2+^ dose-response to ATP is driven by an interplay between the biphasic nature of IP_3_Rs and the desensitization kinetics of P2X7Rs.

**Figure 6.**
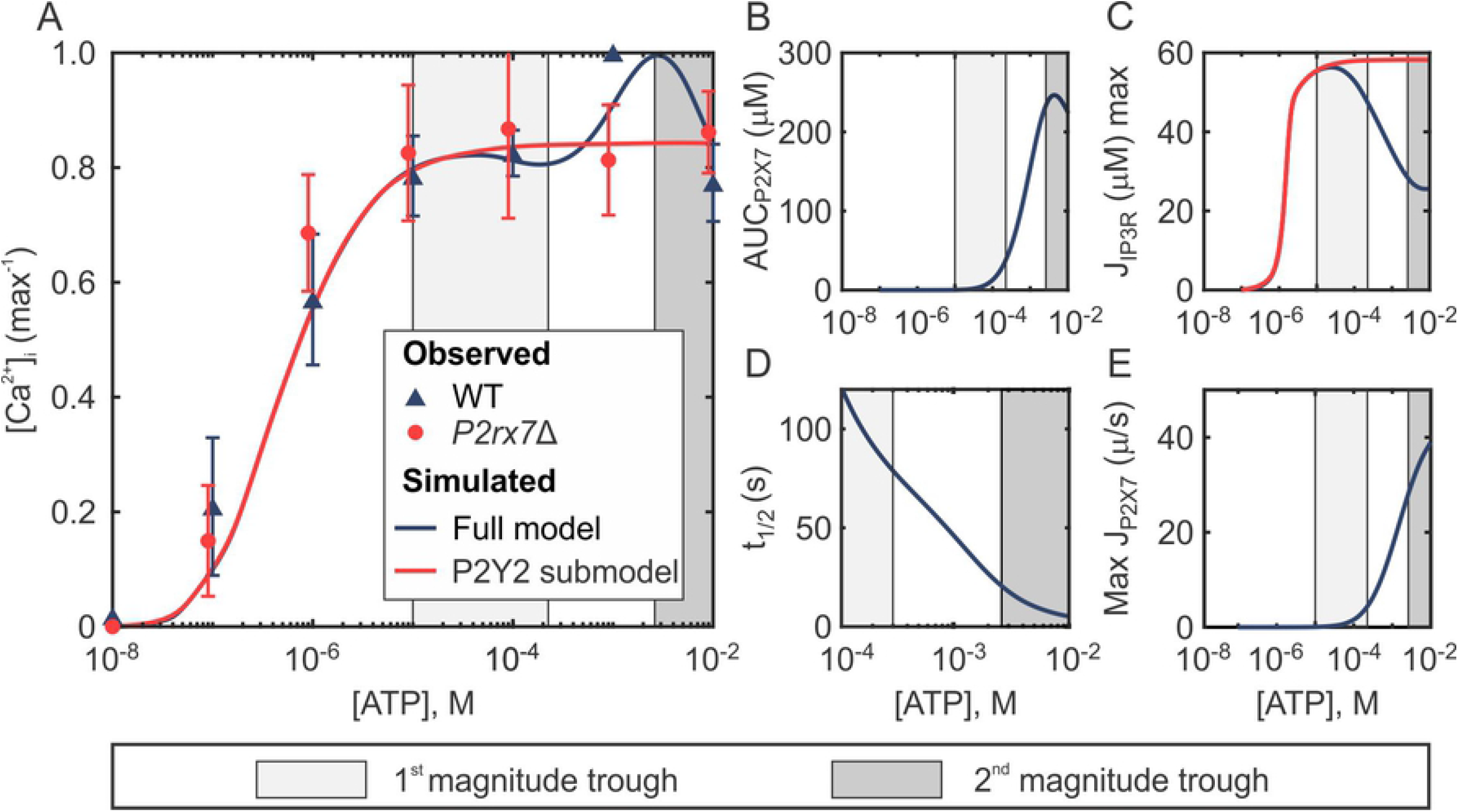
Interaction between P2Y2R and P2X7R underlies the non-monotonic magnitude dose-dependency. **(A)** The magnitude dose-responses of ATP-induced [Ca^2+^]_i_ elevations. Markers indicate experimental means ± SEM in wild-type (WT; *blue*) and *P2rx7*Δ (*red*) C2-OB cells. Curves indicate simulated data generated by the full model (*blue*) and P2Y2 submodel (*red*). **(B)** P2X7R-mediated Ca^2+^ entry estimated from simulated area under the P2X7R flux curve (0-10 s). **(C)** Simulated maximal flux through IP_3_Rs in the full model (*blue*) and P2Y2 only submodel (*red*). **(D)** Rate of P2X7R desensitization estimated from simulated time required for P2X7R flux to decay to half of its maximum. **(E)** Simulated maximum flux through P2X7Rs. *Shaded regions in all panels*: characteristic first (*light grey*) and second (*dark grey*) magnitude troughs observed in WT cells that disappear in *P2rx7*Δ cells.

### 4.6 Contribution of P2Y2 to Ca^2+^ responses at high [ATP]

Next, we examined why the Ca^2+^ response to high [ATP] is dramatically affected in *P2ry2*Δ cells (**Fig. 4B and Fig. 7A**). We examined the magnitudes of Ca^2+^ responses to 10^−2^ M [ATP] in WT *P2ry2*Δ cells, and found that in the absence of P2Y2R, response magnitudes exhibited a distinct bimodal distribution with one cluster of responses similar to those in WT, and another one with much higher response magnitudes (**Fig. 7B**). We hypothesized that the bimodality in the P2X7R-mediated responses is due to P2X7Rs being in the naïve or sensitized initial state (Yan et al., 2010). To verify this, we used the P2X7 submodel to simulate the Ca^2+^ response in two different scenarios: initiating the P2X7-simulations from the naïve closed state *C*_1_ (i.e., *C*_1_(0) = 1, *C*_*i*_(0) = 0, for *i* = 2 ― 4, and *Q*_*i*_(0) = *D*_*i*_(0) = 0, for *i* = 1 ― 4), or from the sensitized closed state *C*_4_ (i.e., *C*_4_(0) = 1, *C*_*i*_(0) = 0, for *i* = 1 ― 3, and *Q*_*i*_(0) = *D*_*i*_(0) = 0, for *i* = 1 ― 4). These two scenarios reflect the heterogeneity in the distribution of unstimulated P2X7R as being predominantly in the naïve or sensitized states. Plotting the [ATP]-dependent dose-response curve for [Ca^2+^]_i_ generated from these time series simulations revealed that P2X7R responses initiated from *C*_1_ produced a dose-response curve that plateaued at a depressed level of [Ca^2+^]_i_ (∼ 40% of max WT response; **Fig. 7A**, *solid red curve*), close to the mean of the left mode of the distribution (**Fig. 7B**). In contrast, P2X7R responses initiated from *C*_4_ state produced a dose-response curve that plateaued at an elevated [Ca^2+^]_i_ (∼ 200% of max WT response; **Fig. 7A**, dashed red curve), closely matching the mean of the right mode of the distribution (**Fig. 7B**). These data suggest that P2Y2 activation may alter the kinetics of P2X7 towards favouring naïve state activation.

**Figure 7.**
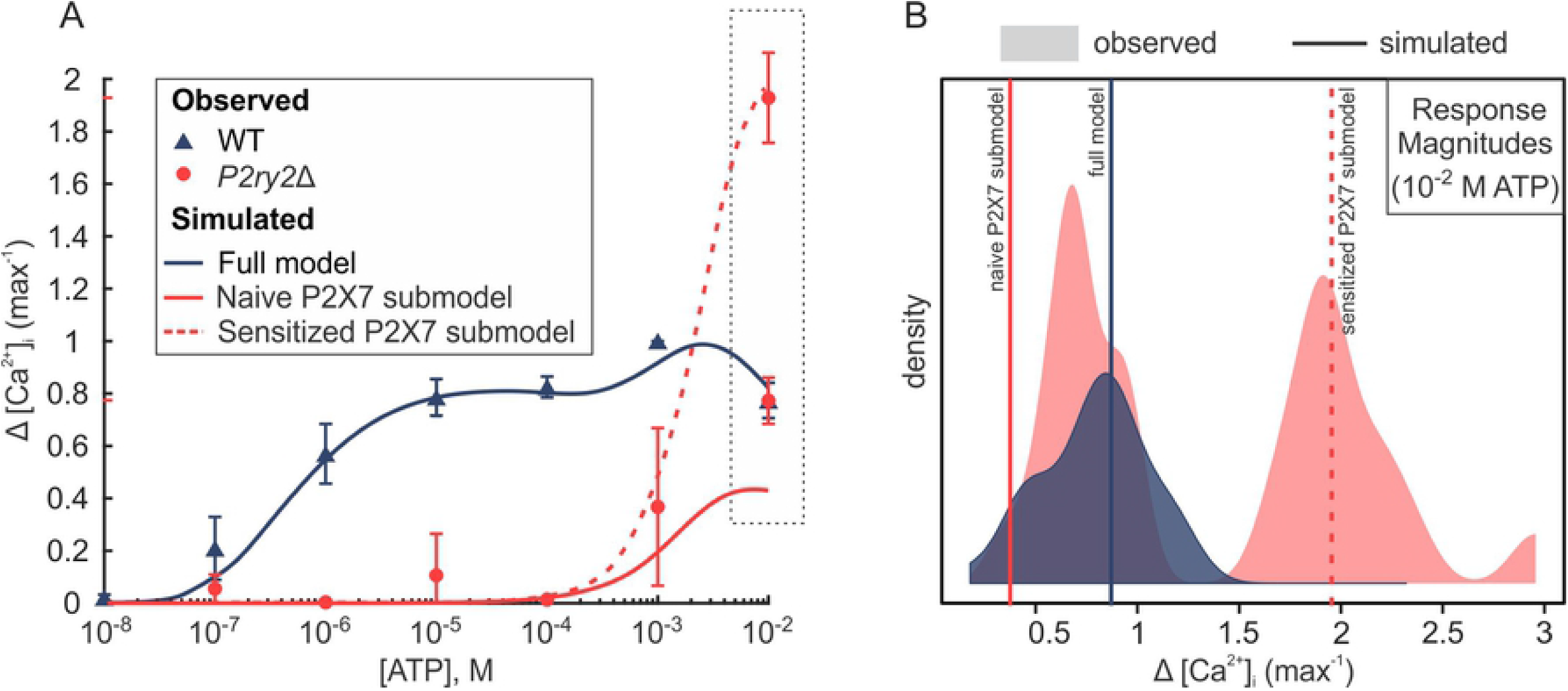
P2Y2 alters P2X7-mediated [Ca^2+^]_i_ response to high [ATP]. **(A)** The magnitude dose-responses of ATP-induced [Ca^2+^]_i_ elevations. Markers indicate experimental means ± SEM in wild-type (WT; *blue*) and *P2ry2*Δ (*red*) C2-OB cells. Curves indicate simulated data generated by the full model (*blue*), a P2X7 submodel initiated from the naïve closed state *C*_1_ (*solid red curve*), or a P2X7 submodel initiated from the sensitized closed state *C*_4_ (*dashed red curve*). **(B)** Density distributions of experimental Ca^2+^ response magnitudes to 10^−2^ M [ATP] in WT cells (*blue density;* unimodal) and *P2ry2*Δ cells (*red density;* bimodal). Vertical lines show simulated response magnitudes (10^−2^ M [ATP]) obtained by the full model (*solid blue line*), naïve P2X7 submodel (*solid red line*) and sensitized P2X7 submodel (*dashed red line*).

### 4.7 Functional contributions of P2Y2 and P2X7 to mechanotransductive signaling

To investigate the potential functional consequences of the complex interactions between P2Y2 and P2X7, we examined how the absence of each of these receptors affects ATP-mediated mechanotransduction. We have previously shown that mechanical stimulation of a single “*primary*” osteoblast with a glass micropipette leads to the release of 10^−5^ to 10^−4^ M ATP into the pericellular space, which then diffuses to stimulate neighbouring non-mechanically perturbed “*secondary*” cells (Mikolajewicz et al., 2019; Mikolajewicz, Zimmermann, et al., 2018). Here, we mechanically stimulated a single fura2-loaded osteoblast from parental C2-OB, or clones deficient in P2Y2, *P2ry2*Δ, or P2X7, *P2rx7*Δ and recorded [Ca^2+^]_i_ responses in the primary and secondary cells (**Fig. 8A**). We found that while in *P2rx7*Δ cells the response was qualitatively similar to WT, in *P2ry2*Δ cells secondary responses were abolished (**Fig. 8B, C**). Quantitatively, the primary response was unaffected in *P2ry2*Δ cells, but exhibited higher magnitude and faster decay in *P2rx7*Δ cells (**Fig. 8C, D**). The suppression of the secondary response in *P2ry2*Δ cells was evident by the reduced signaling radius (p = 9 × 10^―3^), fractions of responding cells (p = 5 × 10^―11^), response magnitudes (p = 2 × 10^―4^) and areas under the curves (AUC; p = 0.02) (**Fig. 8B-D**, *yellow*). In contrast, in *P2rx7*Δ cells, the signaling radius and fraction of responding secondary cells was unaffected; however, the response magnitudes and areas under the curves of secondary cells were significantly higher in *P2rx7*Δ cells compared to parental C2-OB cells (**Fig. 8C, D**, *grey*). These data demonstrate that P2Y2 receptor is critical for the secondary responses, consistent with its high sensitivity to ATP. In addition, the contribution of P2X7 to Ca^2+^ responses is evident even though extracellular ATP in these experiments remained below [ATP] required to stimulate P2X7. Taken together, these data strongly support the importance of an interplay between P2Y2R and P2X7R.

**Figure 8.**
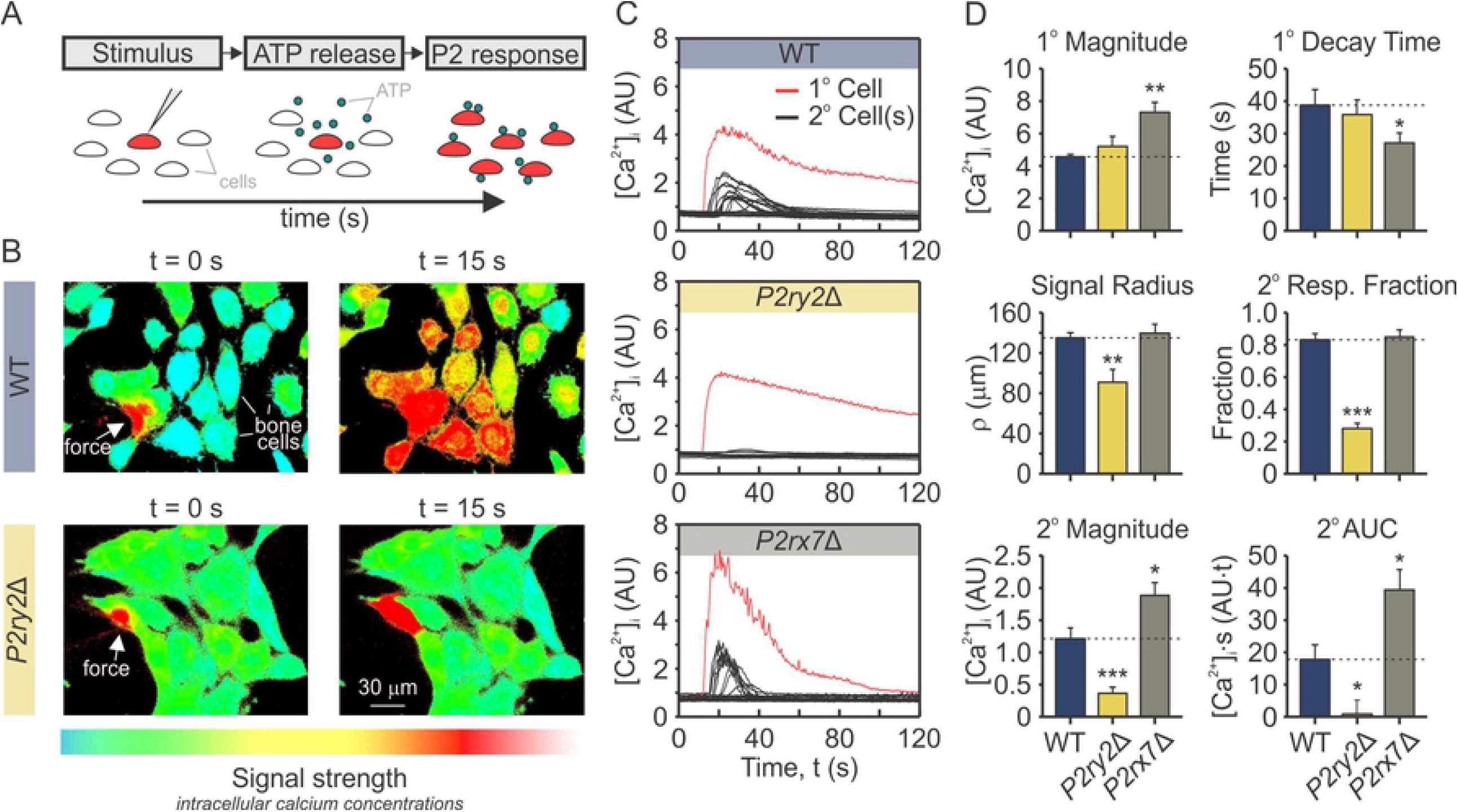
P2Y2R and P2X7R contribution to mechanically stimulated signals in bone cells. **(A)** Fura2-loaded C2-OB cells were plated on a glass-bottom dish and individual cell (*primary; 1°*) was mechanically stimulated with a glass-micropipette inducing ATP release into the extracellular space, which subsequently stimulated P2 responses in neighbouring cells (*secondary; 2°*). **(B)** Representative images of [Ca^2+^]_i_ (pseudocolor of 340/380 ratio) in C2-OB parental (*top*) and *P2ry2*Δ (*bottom*) cultures, in which a single cell was mechanically stimulated at t=0 s (white arrows). The snapshot at 15 s demonstrate secondary responses in neighboring cells. *Red traces*: primary responses; *Black traces*: secondary responses. (**C**) Time-series recordings in WT (*top panel*), *P2ry2*Δ (*middle panel*), and *P2rx7*Δ (*lower panel*) cells. **(D)** Quantification of primary and secondary [Ca^2+^]_i_ response parameters, including signaling radius, fractions of responding cells, response magnitudes and areas under the curves in WT (*blue*), *P2ry2*Δ (*yellow*), and *P2rx7*Δ (*grey*) cells. Data are means ± SEM, *p<0.05, **p<0.01 and ***p<0.001 indicate comparisons between WT and *P2ry2*Δ or *P2rx7*Δ cells assessed by ANOVA and Bonferroni-corrected t-test. AU: arbitrary units.

## 5 Discussion

In this study, we demonstrated that the patterns of P2R-mediated Ca^2+^ responses to ATP are conserved across three independent murine osteoblast models and identified the P2Y2 and P2X7 receptors as the dominant P2 receptors contributing to these responses. We constructed a flux-balance based mathematical model of Ca^2+^ signals induced by the activation of high affinity P2Y2R and low affinity P2X7R. Model predictions were validated by comparing the results of simulations to experimental data of ATP-generated Ca^2+^ signaling in parental and CRISPR/Cas9 -generated P2Y2 and P2Y7 knockouts in osteoblastic C2C12-BMP cells. We demonstrated that activation of P2Y2R by progressively increasing [ATP] induces a transition from transient to oscillatory to transient Ca^2+^ responses due to the biphasic nature of IP3R activation/inactivation kinetics and the interaction of SERCA pumps with IP3Rs. At high [ATP], activation of P2X7R was found to modulate the peak response magnitudes through an interplay between the biphasic nature of IP_3_Rs and the desensitization kinetics of P2X7Rs. Moreover, our study suggests that P2Y2 activity may also alter the kinetics of P2X7 towards favouring naïve state activation. Finally, we demonstrated that functional consequences of lacking P2Y2 or P2X7 are evident beyond the absence of a signal at the expected range of ATP concentrations. Taken together, these findings support a model in which the response to ATP is not a simple superposition of individual P2 receptors, but a complex context-specific functional response build through interactions of multiple P2 receptors.

We have found that murine osteoblastic cells from different sources exhibit similar patterns in their ATP concentration-dependence of Ca^2+^ responses, including non-trivial features such as the transition of response from transient to oscillatory and back to transient when increasing [ATP], and the presence of two troughs in the plateau phase of the magnitude dose-response curve at high [ATP]. These finding suggest that P2 receptors contributing to these responses are also conserved. P2 receptors are ubiquitously expressed in every mammalian cell, with cell- and tissue-type specific patterns of expression (Burnstock, 2007) Consistent with previous reports (Hoebertz, Townsend-Nicholson, Glass, Burnstock, & Arnett, 2000; Orriss et al., 2012), we demonstrated that the pattern of P2 receptors expressed across different murine osteoblast models is consistent at the mRNA level, and identified ATP-sensitive P2X7 and P2Y2 receptors as the dominant P2 receptor subtypes in osteoblasts. In keeping with their important roles in bone, *P2rx7*^-/-^ and *P2ry2*^-/-^ mice have been shown to exhibit severe bone phenotypes, with *P2rx7*^-/-^mice demonstrating significant deficiency in bone mineral density and truncated response to mechano-adaptive loading (Ke et al., 2003; D. Zeng, Yao, & Zhao, 2019), and *P2ry2*^-/-^mice similarly experiencing osteopenia and altered mechanotransducive responses (Y. Xing et al., 2014). In our study, CRISPR-Cas9 double-nickase generated clonal C2-OB cells lacking *P2ry2* or *P2rx7* showed altered responses to sustained ATP stimulation, which translated into significant changes in mechanotransductive [Ca^2+^]_i_ signaling. Thus, P2Y2 and P2X7 receptors play critical roles in mediating the osteoblast response to ATP, particularly in the context of mechanotransducive signaling in bone.

In every osteoblast model, we found that there was a finite range of [ATP] over which oscillatory [Ca^2+^]_i_ response is prevalent. The oscillatory behaviour was abolished in *P2ry2*Δ cells, but preserved in *P2rx7*Δ cells, strongly implicating P2Y2 as a mediator of oscillations. Using the mathematical model of P2Y2-induced changes in [IP_3_] and [Ca^2+^]_ER_ allowed us to examine the mechanism of transition between oscillatory and non-oscillatory (transient) responses. We found that moderate IP_3_ production evokes an oscillatory response because of two factors: *i*) CICR by the IP_3_Rs that exerts negative feedback on the receptors and inhibits them, and *ii*) the interaction of IP_3_Rs with SERCA that pumps Ca^2+^ back into the ER. In contrast, at low [ATP], the Ca^2+^ released by IP_3_Rs is insufficient to feedback and inhibit the receptors, whereas at high [ATP], IP_3_Rs become constitutively activated, making CICR larger than that in the moderate case but no longer able to inactivate the receptors; this produces, as a result, transient Ca^2+^ responses in both cases. While this model prediction is interesting, its validation is limited by difficulties in experimentally measuring osteoblastic IP_3_ dynamics. Specifically, little is known about the basal [IP_3_] (assumed to be 0 μM in our model), which plays a significant role in whether solutions will pass through the oscillatory region obtained in the [IP_3_] and [Ca^2+^]_ER_ space. Furthermore, the ability of the model to predict some of the experimental response profiles observed at elevated [ATP] is limited by the chosen P2Y2R-IP_3_R sub-model, which is unable to slowly decay after a rapid increase in [Ca^2+^]_i_ (Y. Li & Rinzel, 1994). Given the large heterogeneity of responses at these concentrations, choosing a simplified P2Y2R-IP_3_R sub-model was prioritized over the ability to reproduce some response patterns. In spite of these limitations, experimental and modeling data agree on the critical role of P2Y2-induced IP3-mediated Ca^2+^ release from ER in generating oscillatory Ca^2+^ responses to ATP.

Another conserved feature in ATP dose dependence in osteoblasts is the non-monotonic changes in the response magnitude. A similar dose response curve, with troughs at 10^−4^ M and 10^−2^ M ATP and a peak at 10^−3^M ATP was reported in MC3T3-E1 osteoblasts (S. Xing et al., 2016). While we have previously suggested that the decrease in the response magnitude may be mediated by negative effects of one of the receptors that are active at mid-range [ATP] (S. Xing et al., 2016), current study demonstrates that similar regulation is achieved through interactions between P2Y2 and P2X7. In particular, we have found that these characteristic throughs disappear in *P2rx7*Δ cells. Using the mathematical model, we showed that at 10^−4^ M ATP, the additional Ca^2+^ influx through now activated P2X7 inhibits P2Y2-induced IP_3_R activity due to the biphasic dependence of IP_3_Rs on [Ca^2+^]_i_. As a result, the peak [Ca^2+^]_i_ response, which is predominantly mediated by IP_3_R activity, decreases, creating the first trough in the dose response curve at 10^−4^ M [ATP]. With further increase in [ATP], the activation of P2X7Rs, known to monotonically increase with [ATP], starts to outweigh the reduced IP_3_R activity, causing the peak [Ca^2+^]_i_ response to rise, reaching a global maximum around 10^−3^ M ATP. After that, the rate of P2X7R desensitization (that increases with [ATP]) becomes large enough to impede Ca^2+^ influx through the receptors, resulting in a decrease in the peak [Ca^2+^]_i_ response (now predominantly mediated by P2X7Rs) which creates the second trough in the dose response curve at 10^−2^ M ATP. These data demonstrate how activation of low affinity P2X7 may affect the responses mediated by the high affinity P2Y2. Our study also suggests the reciprocal effect of P2Y2 on the function of P2X7 through facilitating the naïve state activation of P2X7. Indeed, previous studies have documented such an effect through the allosteric regulation of P2XR (including P2X7R) by Ca^2+^ (Khadra et al., 2012; Yan et al., 2011), suggesting that Ca^2+^ release from the ER through the P2Y2 pathways may underly the altered kinetics of P2X7. Taken together, our study demonstrates multiple points of interactions between P2Y2 and P2X7 receptors, which are not only activated at very different ranges of ATP concentration, but also belong to different classes of receptors.

Finally, our study demonstrates that the absence of either P2Y2 or P2X7 has significant implications on ATP-mediated mechanotransduction. We used an experimental setup in which the mechanical stimulation of a single osteoblasts generates a micro-injury in its cell membrane, leading to a release of ATP that signals to neighboring (secondary) cells (Mikolajewicz, Zimmermann, et al., 2018). First, we showed that in the absence of P2Y2, the transmission of ATP signal to neighboring cells is effectively interrupted. These findings are consistent with previous reports that osteoblasts from P2Y2^-/-^mice exhibited dramatic reduction in fluid flow-induced Ca^2+^ responses even though the ATP release was similar to WT osteoblasts (Y. Xing et al., 2014). Second, we have found that in the absence of P2X7, both primary and secondary responses are significantly altered. While local ATP concentrations at the site of micro-injury may support the involvement of P2X7 in generating the Ca^2+^ response of the primary cell (Mikolajewicz et al., 2019; Mikolajewicz, Zimmermann, et al., 2018), the observed changes in the secondary responses are surprising, since we have previously shown that the amount of ATP released in these experiments is below the concentrations required for P2X7 activation (Mikolajewicz, Zimmermann, et al., 2018). Nevertheless, this observation is consistent with previously suggested alterations in mechanotransductive signaling in P2X7 deficient mice (Ke et al., 2003; D. Zeng et al., 2019). Thus our study supports the important role of P2 receptor network in generating a mechanotransductive signal that conveys complex information to neighbouring cells

In conclusion, this study provided a complex mechanism of interdependency between the action of high affinity G-protein coupled receptor P2Y2 and a low affinity ligand gated ion channel P2X7. Using a combination of experimental studies in osteoblastic cells with the full compliment of P2 receptors, as well as osteoblasts deficient in P2Y2 or P2X7, and mathematical modeling of P2Y2R-mediated Ca^2+^ release coupled to a Markov model of P2X7R dynamics, allowed us to explore the intricate details of the subcellular signaling induced by ATP in bone forming osteoblasts. The conclusions drawn demonstrated causative links between the exposure to mechanical force, early ATP-mediated signaling, and mechanoadaptive response of bone tissue.

## 6 Materials and Methods

### Software

Figure preparation: CorelDRAWX8 (Corel); Mathematical Modeling: MATLAB R2018a (MathWorks), XPPAUT 8.0. Statistical Analysis: R version 4.0.0 (R Foundation for Statistical Computing);

### Cell Culture

The C2C12 cell line (ATCC CRL-1772) stably transfected with BMP-2 (C2-Ob cells) was plated at 10^4^ cells/cm^2^ in DMEM (supplemented with 10% FBS, 1% sodium pyruvate, 1% penicillin streptomycin) and cultured for 2-3 days prior to experiments. Absence of mycoplasma contamination was verified in cryo-preserved stocks of C2-OB cells using PCR-based detection kit.

### Generation of P2R knockout cell lines

C2-Ob cells were plated in 6-well plates at 100,000 cell/well density 2 days prior to transfection. On the day of transfection, 7.5 µL lipofectamine was diluted in 125 µL Opti-MEM medium (Solution A) and 5 µg *P2ry2* or *P2rx7* CRISPR/Cas9 plasmid and 10 µL P3000 reagent were diluted in a separate 125 µL aliquot of Opti-MEM (Solution B). Solutions A and B were then pooled in a 1:1 ratio and incubated at room temperature for 15 min. Cell media was aspirated and 250 µL of the pooled DNA-lipid complex solution was added to cells and left to incubate at 37 °C for 3 days. 3 days post-transfection, cells were visualized using fluorescent microscope to verify successful transfection through the presence of GFP-positive cells. Transfected cultures were transferred to fresh DMEM media and treated with 5 µM puromycin for 7 days to select for puromycin-resistant clones. After selection, cells were transferred into puromycin-free media, allowed 3 days for recovery, and re-plated in 96 well plates at a ∼1 cell per well density. After 3 weeks of expansion, half of each single-cell colony was re-plated in 96-well plates and the other half was collected for genomic DNA extraction using DNeasy kit. Genomic DNA for each single-cell colony was amplified by touchdown PCR using primer sets designed to flank the genomic region targeted by Cas9 (**Table S2**), and amplicons were separated on a gel to screen for clones with evident band shifts. Selected clones were subsequently validated by immunoblot analysis, and termed *P2ry2*Δ and *P2rx7*Δ cells, for *P2ry2* and *P2rx7*, respectively.

### Quantitative real-time polymerase chain reaction (qRT-PCR)

Total RNA was isolated using RNeasy kit and QIAshredder columns and reverse transcribed using cDNA reverse transcription kit. Real time qPCR was performed using QuantStudio 7 Flex PCR System, with SYBR Green or TaqMan Master Mix. Primer sequences are provided in **Table S2** and cycling conditions in **Table S3**.

### Intracellular Ca^2+^ recordings and analysis

Cells were plated on glass-bottom 35 mm dishes or 48-well plates (MatTek Corporation), for single-cell mechanical stimulation and agonist application experiments, respectively. Cell were loaded with Fura2-AM for 30 min, acclimatized in physiological solution (PS) for 10 min on the stage of an inverted fluorescence microscope (Nikon T2000), and imaged as described previously (Mikolajewicz, Zimmermann, et al., 2018). The Ca^2+^ response parameters were analyzed using a previously developed MATLAB algorithm (https://github.com/NMikolajewicz/Calcium-Signal-Analyzer) (Mackay et al., 2016). To assess ATP dose-dependencies, Fura2-loaded C2-Ob or CB-Ob cells were bathed in 270 µL PS and 30 µL of UDPG, ATP or ADP solutions at 10× final concentration were added (e.g., 30 µL of 10^−5^ M ATP solution was added to cells to achieve 10^−6^ M ATP stimulation).

### Immunoblotting

Cell lysates were extracted in RIPA lysis buffer and samples were prepared and subject to SDS-PAGE on a 10% (w/v) acrylamide gel as previously described (Mikolajewicz, Zimmermann, et al., 2018). Blotted nitrocellulose membranes were incubated with primary antibodies overnight (1:1000 dilution, 5% BSA in TBST, 4°C) and secondary antibodies were applied for 1 h (1:1000 dilution, 5% BSA in TBST, rt) prior to visualization with chemiluminescence system.

### Mechanical-stimulation

Single osteoblastic cells were stimulated by local membrane indentation with a glass micropipette using a FemtoJet microinjector NI2 (Eppendorf Inc.), as previously described (Mikolajewicz, Zimmermann, et al., 2018).

### Statistical Analysis

Data are presented as representative images, means ± standard error (SE) or means ± 95% confidence intervals (CI), as specified in each figure panel along with sample sizes *N* (number of independent experiments) and *n* (number of technical replicates). Curve fitting and [Ca^2+^]_i_ transient characterization were performed in R. Statistical significance was assessed by one- or two-way ANOVA (as specified) and post-hoc two-way unpaired Students t-tests were adjusted using the Bonferroni correction. Significance levels were reported as single symbol (*p<0.05), double symbol (**p<0.01) or triple symbol (***p<0.001).

### Mathematical Model

The mathematical model consisted mainly of **Eqs. (1)-(3)**. The individual terms *J*_*INleak*_, *J*_*IPR*_, *J*_*ERleak*_, *J*_*PMCA*_, *J*_*SERCA*_ and *J*_*P2X7*_, listed in **Eqs. (1)** and **(2)**, were the key Ca^2+^ fluxes considered in this model, as described below.

*(1) Plasma Membrane Ca*^*2+*^ *Leak (J*_*INleak*_). A constant inward leak across cell membrane to ensure that total [Ca^2+^] within the cell remained positive. It was assumed to be constant (see **Table 1**).

*(2) IP*_*3*_*R Ca*^*2+*^ *Flux (J*_*IP3R*_): The Ca^2+^ flux through IP_3_Rs, given by

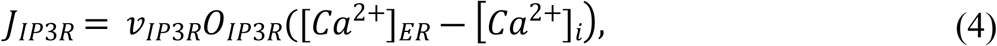

where *v*_*IP3R*_ is the maximum rate of Ca^2+^ release by the IP_3_R and *O*_*IP3R*_ is the IP_3_R open probability, defined by

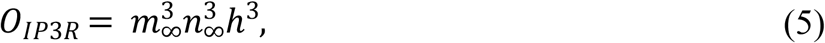

which follows a Hodgkin-Huxley gating formalism adopted by De Young and Keizer (De Young & Keizer, 1992) and later simplified by Li and Rinzel (Y. Li & Rinzel, 1994). In this simplification, the activation by IP_3_ (defined by *m*_∞_) and [Ca^2+^]_i_ (defined by *n*_∞_) through binding to the receptor were assumed to be instantaneous, given by

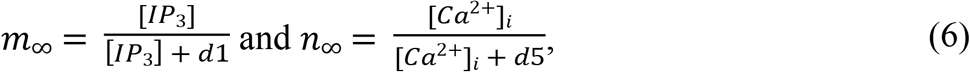

whereas the inactivation by [Ca^2+^]_i_ (defined by the gating variable h), also through binding, was assumed to occur at a much slower time scale governed dynamically by

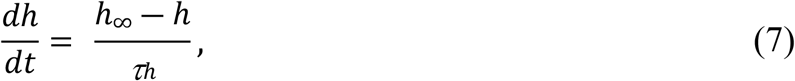

Where

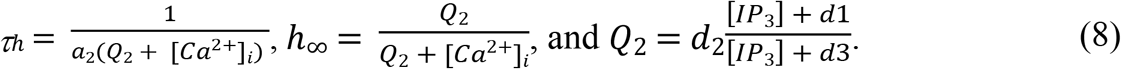

In the study by De Young and Keizer (De Young & Keizer, 1992) the values of *d*_*i*_ (*i* = 1, 2, 3, 5) were fit to experimental data (Watras & Ehrlich, 1991). Note that the dependence of activation and inactivation of *O*_*IP3R*_ on [Ca^2+^]_i_ in Eq. (5) due to CICR makes the profile of IP_3_R open probability biphasic with respect to [Ca^2+^]_i_.

*(3) ER Ca*^*2+*^ *Leak (J*_*ERleak*_). A small leak across the ER membrane (Y. Li & Rinzel, 1994), given by

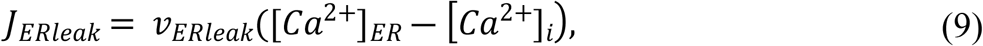

where *v*_*ERleak*_ is the maximal rate of Ca^2+^ leak from the ER.

*(4) Ca*^*2+*^ *ATPase Activity (J*_*PMCA*_ and *J*_*SERCA*_). Ca^2+^ removal by PMCA and Ca^2+^ re-uptake into the ER by SERCA described by Hill functions (Chen et al., 2014; Croisier et al., 2013), given by

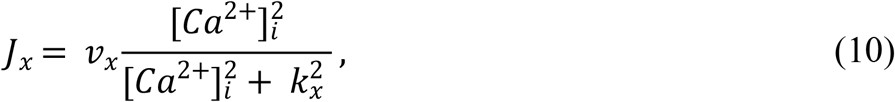

where *x* = *PMCA, SERCA, v*_*x*_ is the maximal pumping rate and *K*_*x*_ is the affinity of the pump to bind to Ca^2+^.

*(5) P2X7R Ca*^*2+*^ *Flux (J*_*P2X7*_). Ca^2+^ flux through P2X7Rs. A 12-state Markov model (Khadra et al., 2013) was initially used to compute the current, given by

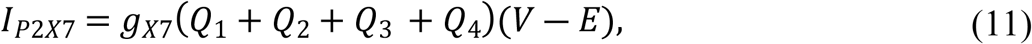

where *g*_*X7*_ is the conductance of the P2X7R open states *Q*_*i*_, *i* = 1–4 (Khadra et al., 2013). With emerging evidence suggesting that P2XRs do not dilate (M. Li, Toombes, Silberberg, & Swartz, 2015; Mackay et al., 2017), the maximum conductance of open (*Q*_1_ and *Q*_2_) and sensitized/primed (*Q*_3_ and *Q*_4_) states in this P2X7R sub-model were assumed to be equal. We also assumed that the rate of desensitization increased with ATP binding (*H*_2(*C*2)_ < *H*_2(*Q*1)_ < *H*_2(*Q*2)_) and that the open probability is higher in the desensitized row (i.e., *K*_7_ > *K*_*j*_, *j* = 2, 4, 6). These modifications kept the time series simulations of the P2X7R model generally unchanged. To obtain the overall Ca^2+^ flux through these channels, we then used the formalism from Zeng et al. (S. Zeng, Li, Zeng, & Chen, 2009) to convert ionic current to flux, scaled by a fraction that represents the average Ca^2+^ flux (Egan & Khakh, 2004) relative to that for Na^+^ and K^+^. The latter was necessary as P2X7Rs are non-selective cation channels. Using the above description, the following expression was used to describe this flux

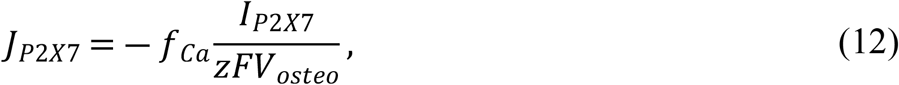

where *z* is the valence of Ca^2+^ (*z* = +2), *F* is Faraday’s constant, *V*_*osteo*_ is the volume of the osteoblast in liters and *f*_*Ca*_ is the fraction of Ca^2+^ flux through P2X7R.

### Software and numerical methods

All time-series simulations were performed in MATLAB (Mathworks, Natick, MA). Initially, all simulations were run for a period of 2000 s in the absence of ATP to obtain the resting state of the cell. The basal [IP_3_] (the [IP_3_] in the absence of extracellular ATP) was assumed to be zero. P2Y2R and P2X7R knockout recordings were simulated by setting *J*_*P2X7*_ and *J*_*IP3R*_ (**Eqs. (5) and (12)**) to zero, respectively. These simulations were then used to compute [ATP]-dependent dose-response curves of [Ca^2+^]_i_ by evaluating the maximum Ca^2+^ response at each ATP dose in MATLAB. The bifurcation analysis (**Fig. 5)** was performed using XPPAUT (a freeware available online at **http://www.math.pitt.edu/~bard/xpp/xpp.html**). To facilitate reproduction of results, the codes used to perform simulations of the model can be obtained online (Mikolajewicz, Smith, Komarova, & Khadra, 2021). These simulations can be run by solving the function “fullmodel.m” using the ordinary differential equation solver ode15s. Figure 4A, Figure 6 and Figure 7 can be obtained by running the files titled “Figure4A.m”, “Figure6.m” and “Figure7.m”, respectively.

## 8 Author Contributions

Study conception and design: NM, SVK, AK

Acquisition of experimental data: NM

Mathematical modelling: DS, AK

Analysis and interpretation of data: NM, DS, SVK, AK

Drafting of Manuscript: NM, DS

All authors contributed to the critical revision and approval of the final manuscript.

## 9 Acknowledgements

This work was supported by the Natural Sciences and Engineering Research Council of Canada discovery grants to AK and SVK, the *Fonds Nature et technologies* team grant to AK and the Canadian Institutes for Health Research (CIHR MOP-77643) grant to SVK. NM was supported by the Faculty of Dentistry, McGill University, and le Réseau de Recherche en Santé Buccodentaire et Osseuse (RSBO). Special thanks to Dr. M. Murshed (McGill, Montreal) for C2-OB cells and Dr. P. Grutter and his graduate students M. Anthonisen and M. Rigby (McGill, Montreal) for glass capillary puller.

## 11 Supporting Information

### Solutions and Reagents

#### Solutions

Phosphate-buffered saline (PBS; 140 mM NaCl, 3 mM KCl, 10 mM Na_2_HPO_4_, 2 mM KH_2_PO_4_, pH 7.4), autoclaved; Phosphate buffered saline with Tween 20 (PBST; PBS + 1% Tween 20); Physiological solution (PS; 130 mM NaCl, 5 mM KCl, 1 mM MgCl_2_, 1 mM CaCl_2_, 10 mM glucose, 20 mM HEPES, pH 7.6), sterilized by 0.2 µm filtration; RIPA lysis buffer (50 mM Tris, pH 7.4, 150 mM NaCl, 1% Nonidet P-40, 1 mM EDTA, 1 mg/mL aprotinin, 2 mg/mL leupeptin, 0.1 mM phenylmethylsulfonyl fluoride, 20 mM NaF, 0.5 mM Na3VO4); TBST buffer (10 mM Tris-HCL, pH 7.5, 150 mM NaCl, 1% Tween 20).

#### Reagents

High-capacity cDNA reverse transcription kit (Cat. 4368814); *Power* SYBR Green Master Mix (Cat. 4368702) from Applied Biosystems. Nitrocellulose membrane, 0.45 µm (Cat. 162-0115) from Bio-Rad. Opti-MEM (Cat. 31985062) from Gibco. Fura2-AM (Cat. F1221); Lipofectamine 3000 transfection reagent (Cat. L3000001); Quant-iT protein assay kit (Cat. Q33210) from Invitrogen. Puromycin (Cat. ant-pr-1) from InvivoGen. 35 mm glass-bottom dishes (Cat. P35G-1.5-14-C); 48-well glass-bottom plates (Cat. P48G-1.5-6-F) from MatTek Corporation. RNeasy Mini Kit (Cat. 74104) from Qiagen. Adenosine 5’-triphosphate (ATP; Cat. A9187); Venor GeM Mycoplasma PCR-based detection kit (Cat. MP0025) from Sigma-Aldrich. Pierce ECL western blotting substrate (Cat. 32106) from Thermo Scientific. Dulbecco’s modified eagle medium (DMEM; Cat. 319-020 CL); Fetal bovine serum (FBS; Cat. 080152); Penicillin streptomycin (Cat. 450-201-EL); Sodium pyruvate (Cat. 600-110-UL) from Wisent Bio Products.

### 11.1 Supplemental Figures

**Figure S1. Aligning and scaling ATP dose-dependence curves.** (**A-C**) Schematic illustrating processing of ATP-dose-dependent response curves (**A**). ATP dose-dependent responses from three independent murine cell lines (**A**, *left panel;* **B**) were aligned using a linear transformation to match peaks/troughs (**A**, *middle panel;* **C**) and responses were rescaled to [0,1] interval (**A**, *right panel;* **Fig. 1C**). *Curves*: Loess curves; *Markers*: Response means (*circle*: BM-OB; *triangle*: C2-OB; *square*: CB-OB).

### 11.2 Supplemental Tables

**Supplemental Table 1.** Calcium response parameters.

| Supplemental Table 1. Calcium response parameters. |  |  |  |
| --- | --- | --- | --- |
| Parameter | Symbol | Units | Description |
| Activation time | $t_{10-90\%}$ | s | Time for signal to increase from 10 to 90% of peak. |
| Magnitude | - | $\Delta[\text{Ca}^{2+}]_i$ | Amplitude of response (peak – baseline). |
| Area under curve | $AUC$ | $\Delta[\text{Ca}^{2+}]_i \cdot \text{s}$ | Area under response curve. |
| Decay time | $\tau_{decay}$ | s | Decay constant of the exponential decay region of the deactivation phase of response. |
| Oscillatory fraction | - | - | Fraction of cells with multi-peaked (oscillatory response). |
| Oscillatory magnitude | $E$ | $\Delta[\text{Ca}^{2+}]_i$ | Mean magnitude of oscillatory peaks. |
| Oscillatory period | $T$ | s | Mean periodicity of oscillatory peaks. |
| Oscillatory peaks | $N_{osc}$ | - | Number of observed oscillatory peaks. |
| Response fraction | - | - | Fraction of cells with discernable calcium response. |
| Signal radius | $\rho$ | $\mu\text{m}$ | Average distance of secondary ($2^\circ$) responders from primary ($1^\circ$) stimulated cell. |
| Refer to (Mackay et al., 2016) for schematic-based definitions. |  |  |  |

**Supplemental Table 2.**
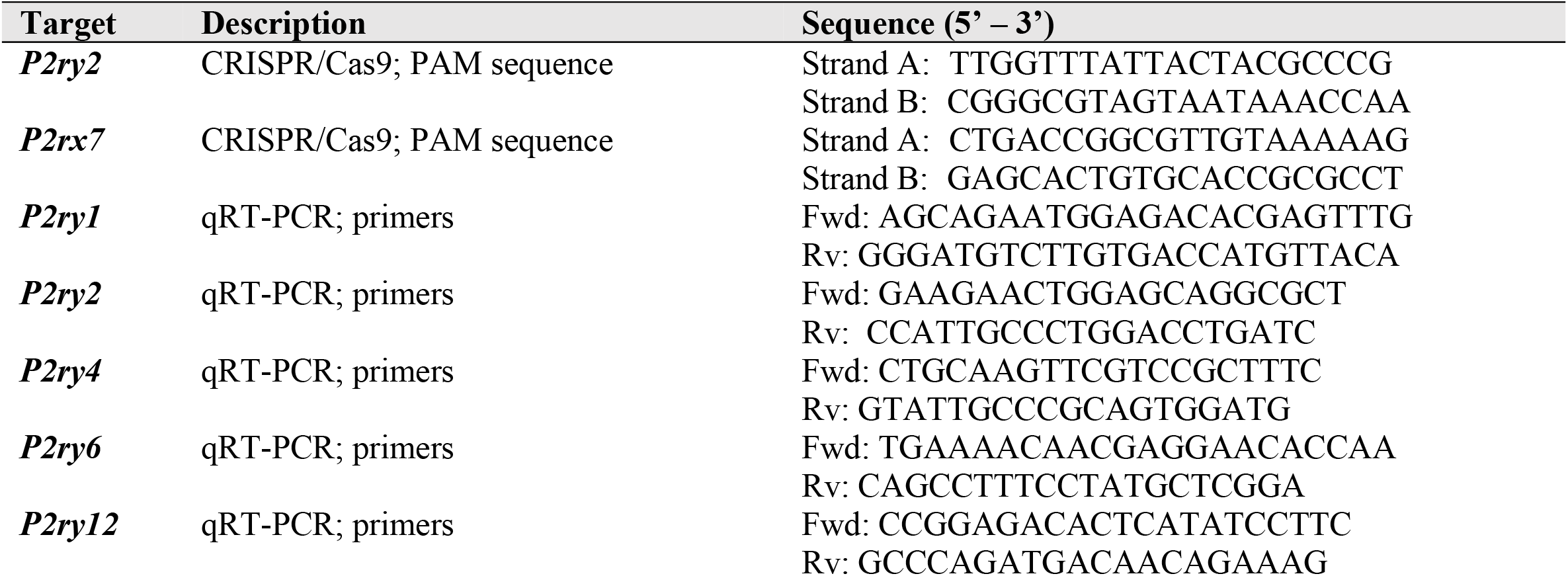
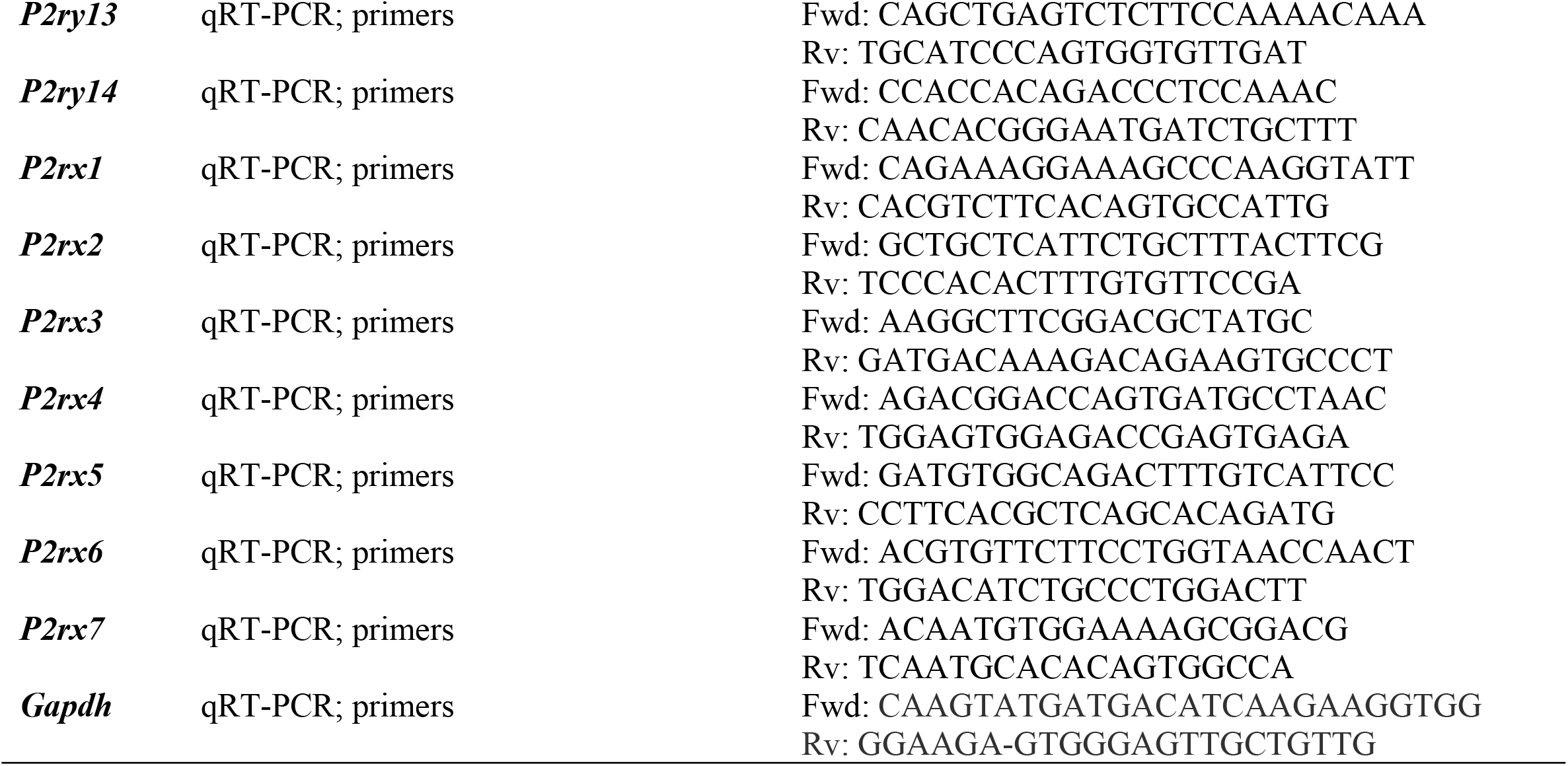
Primer and PAM sequences.

**Supplemental Table 3.** PCR cycling conditions.

| Supplemental Table 3. PCR cycling conditions |  |  |
| --- | --- | --- |
| Stage | Description | Cycles (n) |
| **qRT-PCR (cDNA template)** |  |  |
| **Phase 1** | Denaturation: 94 °C, 180 s | 1 |
| **Phase 2** | Denaturation 94 °C, 30 s<br>Annealing: 60 °C, 45 s<br>Elongation: 72 °C, 60 s | 40 |
| **Phase 3** | Elongation: 72 °C, 600 s<br>Hold: 4 °C | 1 |

## References

Bours, M., Dagnelie, P. C., Giuliani, A. L., Wesselius, A., & Di Virgilio, F. (2011). P2 receptors and extracellular ATP: a novel homeostatic pathway in inflammation. Front Biosci (Schol Ed), 3, 1443–1456.

Burnstock, G. (2007). Physiology and pathophysiology of purinergic neurotransmission. Physiological reviews.

Burnstock, G. (2018). Purine and purinergic receptors. Brain and Neuroscience Advances, 2, 2398212818817494.

Burnstock, G., & Ralevic, V. (2014). Purinergic signaling and blood vessels in health and disease. Pharmacological reviews, 66(1), 102–192.

Chen, Y.-f., Cao, J., Zhong, J.-n., Chen, X., Cheng, M., Yang, J., & Gao, Y.-d. (2014). Plasma membrane Ca2+-ATPase regulates Ca2+ signaling and the proliferation of airway smooth muscle cells. European Journal of Pharmacology, 740, 733–741. doi:https://doi.org/10.1016/j.ejphar.2014.05.055

Coddou, C., Yan, Z., Obsil, T., Huidobro-Toro, J. P., & Stojilkovic, S. S. (2011). Activation and regulation of purinergic P2X receptor channels. Pharmacological reviews, 63(3), 641–683.

Croisier, H., Tan, X., Perez-Zoghbi, J. F., Sanderson, M. J., Sneyd, J., & Brook, B. S. (2013). Activation of store-operated calcium entry in airway smooth muscle cells: insight from a mathematical model. PLoS One, 8(7), e69598.

De Young, G. W., & Keizer, J. (1992). A single-pool inositol 1, 4, 5-trisphosphate-receptor-based model for agonist-stimulated oscillations in Ca2+ concentration. Proceedings of the National Academy of Sciences, 89(20), 9895–9899.

Egan, T. M., & Khakh, B. S. (2004). Contribution of calcium ions to P2X channel responses. Journal of Neuroscience, 24(13), 3413–3420.

Erb, L., & Weisman, G. A. (2012). Coupling of P2Y receptors to G proteins and other signaling pathways. Wiley Interdisciplinary Reviews: Membrane Transport and Signaling, 1(6), 789–803.

Fedorov, I. V., Rogachevskaja, O. A., & Kolesnikov, S. S. (2007). Modeling P2Y receptor-Ca2+ response coupling in taste cells. Biochim Biophys Acta, 1768(7), 1727–1740. doi:10.1016/j.bbamem.2007.04.002

Grol, M. W., Pereverzev, A., Sims, S. M., & Dixon, S. J. (2013). P2 receptor networks regulate signaling duration over a wide dynamic range of ATP concentrations. J Cell Sci, 126(Pt 16), 3615–3626. doi:10.1242/jcs.122705

Hoebertz, A., Townsend-Nicholson, A., Glass, R., Burnstock, G., & Arnett, T. (2000). Expression of P2 receptors in bone and cultured bone cells. Bone, 27(4), 503–510.

Jacobson, K. A., Costanzi, S., Joshi, B. V., Besada, P., Shin, D. H., Ko, H., … Mamedova, L. (2006). Agonists and antagonists for P2 receptors. Paper presented at the Novartis Foundation symposium.

Ke, H. Z., Qi, H., Weidema, A. F., Zhang, Q., Panupinthu, N., Crawford, D. T., … Audoly, L. P. (2003). Deletion of the P2X7 nucleotide receptor reveals its regulatory roles in bone formation and resorption. Molecular Endocrinology, 17(7), 1356–1367.

Khadra, A., Tomić, M., Yan, Z., Zemkova, H., Sherman, A., & Stojilkovic, S. S. (2013). Dual gating mechanism and function of P2X7 receptor channels. Biophys J, 104(12), 2612–2621. doi:10.1016/j.bpj.2013.05.006

Khadra, A., Yan, Z., Coddou, C., Tomić, M., Sherman, A., & Stojilkovic, S. S. (2012). Gating properties of the P2X2a and P2X2b receptor channels: experiments and mathematical modeling. J Gen Physiol, 139(5), 333–348. doi:10.1085/jgp.201110716

Lemon, G., Brockhausen, J., Li, G. H., Gibson, W. G., & Bennett, M. R. (2005). Calcium mobilization and spontaneous transient outward current characteristics upon agonist activation of P2Y2 receptors in smooth muscle cells. Biophys J, 88(3), 1507–1523. doi:10.1529/biophysj.104.043976

Li, M., Toombes, G. E., Silberberg, S. D., & Swartz, K. J. (2015). Physical basis of apparent pore dilation of ATP-activated P2X receptor channels. Nature neuroscience, 18(11), 1577–1583.

Li, Y., & Rinzel, J. (1994). Equations for InsP3 receptor-mediated [Ca2+] i oscillations derived from a detailed kinetic model: a Hodgkin-Huxley like formalism. Journal of theoretical Biology, 166(4), 461–473.

Mackay, L., Mikolajewicz, N., Komarova, S. V., & Khadra, A. (2016). Systematic characterization of dynamic parameters of intracellular calcium signals. Frontiers in physiology, 7, 525.

Mackay, L., Zemkova, H., Stojilkovic, S. S., Sherman, A., & Khadra, A. (2017). Deciphering the regulation of P2X4 receptor channel gating by ivermectin using Markov models. PLoS computational biology, 13(7), e1005643.

Mikolajewicz, N., Mohammed, A., Morris, M., & Komarova, S. V. (2018). Mechanically stimulated ATP release from mammalian cells: systematic review and meta-analysis. Journal of Cell Science, 131(22).

Mikolajewicz, N., Sehayek, S., Wiseman, P. W., & Komarova, S. V. (2019). Transmission of mechanical information by purinergic signaling. Biophysical journal, 116(10), 2009–2022.

Mikolajewicz, N., Smith, D., Komarova, S. V., & Khadra, A. (2021). High-affinity P2Y2 and low-affinity P2X7 receptor interaction modulates ATP-mediated calcium signalling in murine osteoblasts. Anmar Khadra Repository 2020, Available from: www.medicine.mcgill.ca/physio/khadralab/Codes/code_ploscomp2.html.

Mikolajewicz, N., Zimmermann, E. A., Willie, B. M., & Komarova, S. V. (2018). Mechanically stimulated ATP release from murine bone cells is regulated by a balance of injury and repair. ELife, 7, e37812.

North, R. A. (2002). Molecular physiology of P2X receptors. Physiological reviews.

Orriss, I. R., Key, M. L., Brandao-Burch, A., Patel, J. J., Burnstock, G., & Arnett, T. R. (2012). The regulation of osteoblast function and bone mineralisation by extracellular nucleotides: The role of p2x receptors. Bone, 51(3), 389–400.

Verkhratsky, A., & Burnstock, G. (2014). Biology of purinergic signalling: its ancient evolutionary roots, its omnipresence and its multiple functional significance. Bioessays, 36(7), 697–705.

Watras, J., & Ehrlich, B. E. (1991). Bell-shaped calcium-response curves of lns (l, 4, 5) P 3-and calcium-gated channels from endoplasmic reticulum of cerebellum. Nature, 351(6329), 751–754.

Xing, S., Grol, M. W., Grutter, P. H., Dixon, S. J., & Komarova, S. V. (2016). Modeling interactions among individual P2 receptors to explain complex response patterns over a wide range of ATP concentrations. Frontiers in physiology, 7, 294.

Xing, Y., Gu, Y., Bresnahan, J. J., Paul, E. M., Donahue, H. J., & You, J. (2014). The roles of P2Y 2 purinergic receptors in osteoblasts and mechanotransduction. PLoS One, 9(9), e108417.

Yan, Z., Khadra, A., Li, S., Tomic, M., Sherman, A., & Stojilkovic, S. S. (2010). Experimental characterization and mathematical modeling of P2X7 receptor channel gating. J Neurosci, 30(42), 14213–14224. doi:10.1523/jneurosci.2390-10.2010

Yan, Z., Khadra, A., Sherman, A., & Stojilkovic, S. S. (2011). Calcium-dependent block of P2X7 receptor channel function is allosteric. J Gen Physiol, 138(4), 437–452. doi:10.1085/jgp.201110647

Zemkova, H., Khadra, A., Rokic, M. B., Tvrdonova, V., Sherman, A., & Stojilkovic, S. S. (2015). Allosteric regulation of the P2X4 receptor channel pore dilation. Pflugers Arch, 467(4), 713–726. doi:10.1007/s00424-014-1546-7

Zeng, D., Yao, P., & Zhao, H. (2019). P2X7, a critical regulator and potential target for bone and joint diseases. Journal of cellular physiology, 234(3), 2095–2103.

Zeng, S., Li, B., Zeng, S., & Chen, S. (2009). Simulation of spontaneous Ca2+ oscillations in astrocytes mediated by voltage-gated calcium channels. Biophysical journal, 97(9), 2429–2437.

